# Sparse cortical dynamics reveal flexible condition-dependent spike-order codes

**DOI:** 10.64898/2026.05.13.724761

**Authors:** Guozhang Chen, Wolfgang Maass, Franz Scherr

## Abstract

Cortical networks compute with remarkably sparse spiking activity, yet the circuit mechanisms that organize these few spikes into flexible, condition-dependent temporal codes remain poorly understood. Here we combine analyses of large-scale mouse recordings with a data-driven cortical microcircuit model (CMM) of mouse V1. In recordings, V1 activity exhibits condition-dependent spike-order sequences: peak-latency order varies with task outcome, current image identity, and preceding image identity, while remaining stable under split-half and single-trial analyses. After task optimization by backpropagation through time, the CMM reproduces this sequence-level signature and sparse activity more closely than the matched randomly connected RSNN and rate-RNN controls tested here. Ablations indicate that neuronal heterogeneity and distance-dependent local connectivity each reduce rigid sequential activity, with their combination giving the closest match to measured cortical signatures. Low-dimensional trajectory visualizations and model-silencing experiments further identify high-mutual-information early neurons whose removal perturbs task trajectories and decisions. Together, these results identify a biologically grounded computational principle: neuronal diversity and local connectivity help sparse recurrent networks avoid rigid temporal pipelines and support flexible, condition-dependent spike-order computation, providing candidate design principles for SNNs that exploit flexible temporal codes.

## 1 Introduction

Sparse spiking activity is a central feature of cortical computation and a major motivation for spiking neural networks (SNNs). Event-driven artificial SNNs can solve demanding tasks, yet task accuracy alone does not guarantee that their internal computations resemble the sparse, temporally structured, and trial-variable dynamics observed in cortex [Olshausen and Field, 1996, Zador, 2019, Zenke and Neftci, 2021, Bellec et al., 2020]. The unresolved question addressed here is therefore narrower and more mechanistic: which circuit constraints allow sparse spiking networks to compute through condition-dependent temporal activity patterns, and which of these constraints can be transferred to artificial SNNs?

This question builds on, rather than replaces, prior work on sequential population coding and recurrent-network dynamics. Sequential and condition-dependent neural activity has been observed in several cortical tasks and areas [Driscoll et al., 2017, Koay et al., 2022, Yiling et al., 2023], and trained recurrent networks can implement flexible computations through task-dependent dynamical motifs [Driscoll et al., 2024]. The remaining gap is to connect such temporal signatures to concrete circuit constraints, for example neuronal heterogeneity and local wiring, in a model that can be systematically ablated.

We address this gap in two steps (Fig. 1A). First, we analyze large-scale *in vivo* recordings from awake mice performing a visual-change-detection task (Fig. 1B) [Garrett et al., 2020, Allen Institute MindScope Program, 2022] and evidence accumulation task (Fig. 1C) [Koay et al., 2022], and use them to define a biological target: sparse, condition-dependent spike-order structure in V1. Second, we test whether a data-driven cortical microcircuit model (CMM) of mouse V1 [Billeh et al., 2020, Chen et al., 2022, Ito et al., 2026], after task optimization, can reproduce this target more closely than matched recurrent controls.

**Figure 1:**
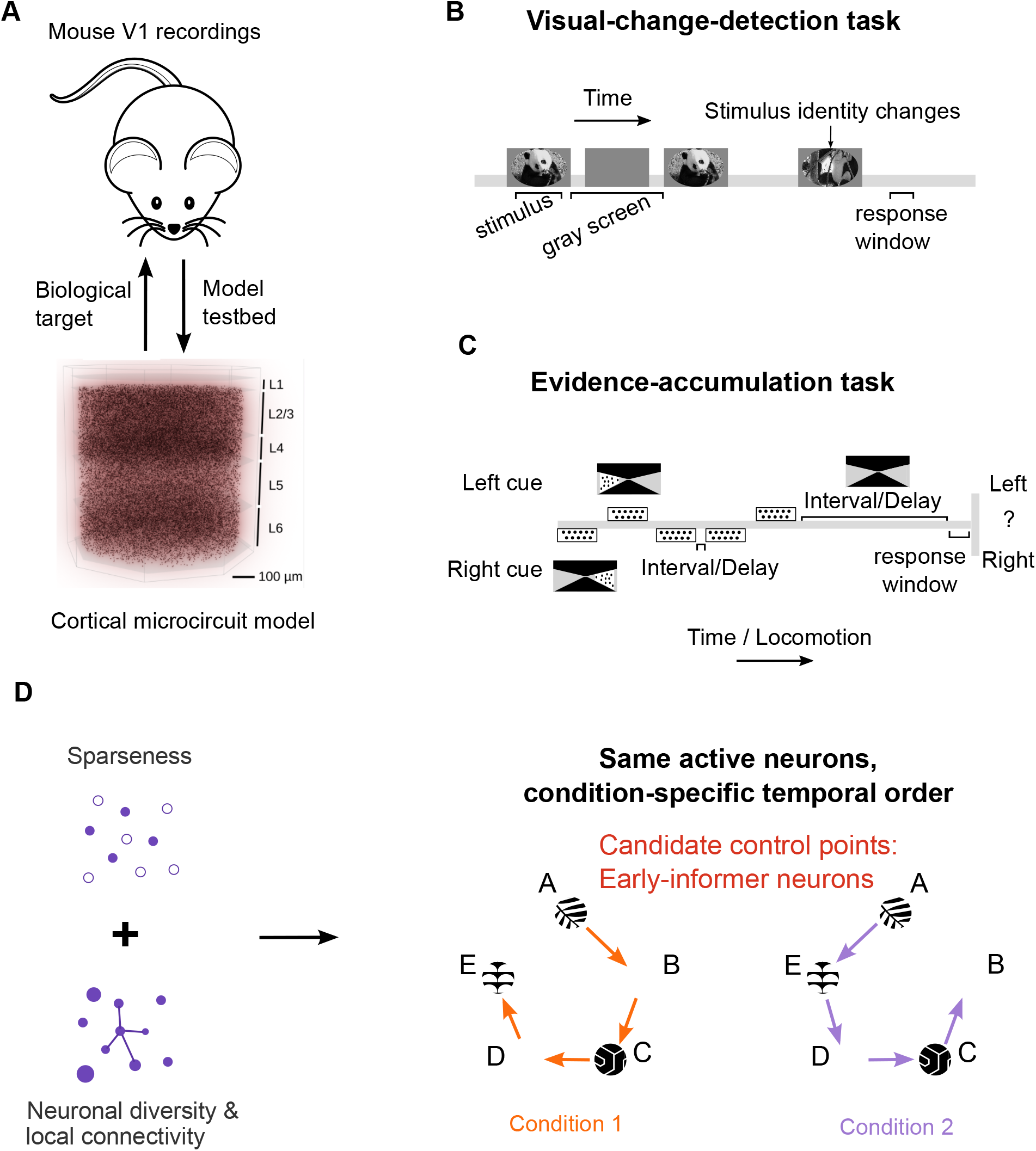
Conceptual overview of sparse, condition-dependent spike-order computation. **(A)** Mouse V1 recordings define the biological target, and a data-driven cortical microcircuit model (CMM) serves as a model testbed for circuit-level analysis. **(B)** In the visual-change-detection task, animals compare the current natural image with the preceding image and report whether the stimulus identity has changed. **(C)** In the evidenceaccumulation task, sequential left and right cues are integrated to determine a later choice. **(D)** Proposed design principle. Sparseness establishes an economical spiking regime, while neuronal diversity and local connectivity structure this activity into flexible, condition-dependent temporal orders. The state-machine schematic illustrates the operational definition of a spike-order code: the same active neurons can be traversed in different spiking orders under different task conditions. Model perturbations further motivate the prediction that early-informer neurons can serve as candidate control points for subsequent trajectory selection and behavioral decisions.

The data and model together support a specific computational hypothesis: sparse cortical activity can support task computation through condition-dependent relative firing order rather than a single static sequence (Fig. 1D). Also, cortical cell-type diversity and distance-dependent local connectivity can reduce the rigid sequential dynamics of generic recurrent networks, facilitating condition-dependent routing that coexists with training-induced sparseness. We further identify high-mutual-information “early-informer” neurons in the trained CMM whose silencing perturbs model trajectories and decisions, suggesting experimentally testable predictions.

## 2 Results

### 2.1 Condition-dependent spike-order structure in mouse V1

To establish a biological baseline for sparse computation, we first examined the dynamics and neural coding of cortical spiking activity during a demanding cognitive task: the visual-change-detection task (Fig. 1B). In this task, mice received a continuous sequence of natural images, separated by delay periods where the screen was empty. The task required the subject to decide whether the currently presented image matched the immediately preceding one. This is a computationally demanding task, as it requires rapid sensory processing, feature extraction, working memory, and comparison.

Mice were trained to solve this task, and large-scale neuron recordings from the primary visual cortex (V1) during task execution were made publicly available [Allen Institute MindScope Program, 2022] (Fig. 1B). We analyzed these recordings and found that V1 activity exhibited sparse, condition-dependent temporal organization. Most V1 neurons fired only within brief, specific time windows that varied across the population, creating a characteristic temporal sequence of distributed sparse activity (Fig. 2A, left and right).

**Figure 2:**
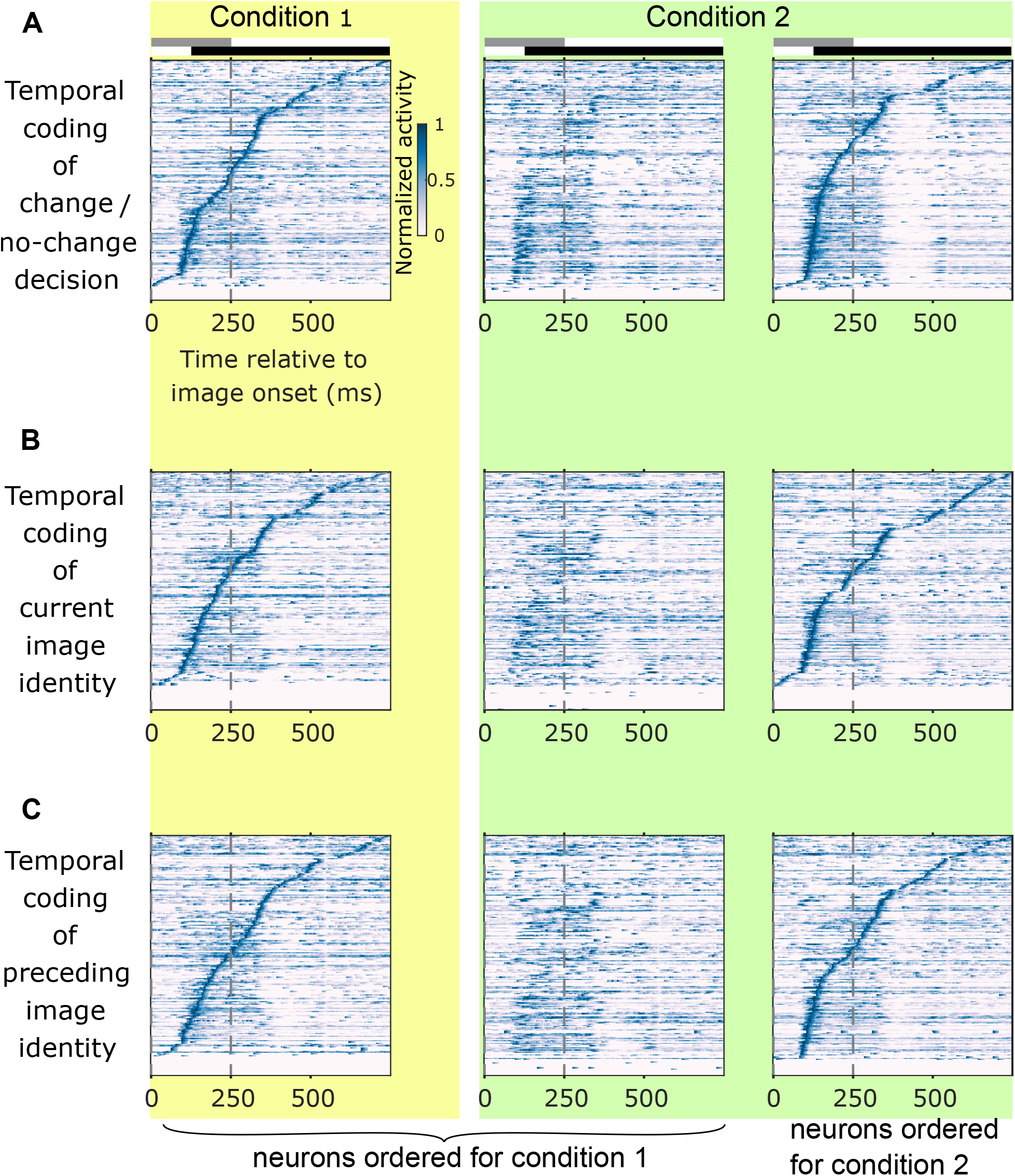
Condition-dependent spike-order structure in mouse V1 during the visual-change-detection task. Each plot shows the normalized average firing activity of 296 neurons recorded from mouse primary visual cortex (V1) during a visual-change-detection task. Activities are averaged across 50 trials and temporally smoothed using a sliding exponential filter (window length: 10 ms, time constant: 20 ms). Each row represents a single neuron, and the blue-to-white color indicates its normalized activity. Neurons (vertical axis) are sorted by the time of their peak activity relative to the stimulus onset (horizontal axis at 0 ms) for condition 1 or 2 (specified at the bottom). The gray bar indicates the image presentation window. The black bar indicates the response window for reporting an image change. The three-column structure in each panel is designed to visually demonstrate that neural firing sequences are different across conditions. **(A)** Comparison of firing order for change and no-change trials. Left: Activity during change trials, with neurons sorted by their peak firing time in this condition, revealing a distinct temporal sequence. Middle: Activity during no-change trials, plotted with the same neuron sorting as in the change condition on the left. The resulting blurred pattern demonstrates that the firing order is significantly different between the two conditions. Right: Activity during no-change trials, now sorted by its own optimal firing order, revealing its own unique and clear temporal sequence. **(B)** Comparison of firing order for two different current images (Image 1 vs. Image 2) which are randomly chosen, while the preceding image and trial outcome (change) are held constant. Condition 1 and 2 are defined by these 2 images, respectively. The blurred middle panel, which shows the activity for Image 2 sorted by the order of Image 1, demonstrates that the neural sequence robustly encodes the identity of the current stimulus. **(C)** Comparison of sequences for two different preceding images (Image 1 vs. Image 2) which are randomly chosen, while the current image and trial outcome (change) are held constant. Condition 1 and 2 are defined by these 2 images, respectively. Middle panel like in B.

Crucially, this sequential organization was highly condition-dependent: the relative firing order of the neurons can distinguish the result of the network computation (change or no-change), as well as the identities of the current and preceding images (Fig. 2B, C). When neurons were sorted by their peak firing times for one specific task outcome (e.g., ‘change’ trials), this clear sequential ordering vanished when the same neurons were plotted during a different outcome (e.g., ‘no-change’ trials) (Fig. 2A, middle column). However, the network exhibited a distinct sequential order associated with the second condition (Fig. 2A, right column). To verify that this observed condition-dependent routing was not a sorting artifact, we confirmed sequence stability using split-half cross-validation (*ρ* = 0.52, *t*-test: *p* < 10^−10^, Methods Sec. 4.5).

These results demonstrate that the V1 activity is not described by a single condition-invariant firing order. Instead, V1 may dynamically route information through flexible, condition-dependent sparse pathways, where the relative firing order encodes a substantial amount of condition-specific information. This biological signature is consistent with findings in other cortical areas and tasks, e.g., in [Driscoll et al., 2017], Fig. 2 of [Koay et al., 2022], and [Yiling et al., 2023], and provides a clear target for evaluating artificial SNN architectures. Throughout the manuscript, we use the term “pathway” operationally to denote condition-specific temporal organization of active neuronal populations; this analysis does not by itself establish a traced anatomical pathway or a causal biological routing mechanism.

### 2.2 A trained cortical microcircuit model reproduces the cortical spike-order signature

To test which architectural constraints can support the condition-dependent spike-order structure described above, we used a data-driven cortical microcircuit model (CMM) of mouse V1 [Billeh et al., 2020, Chen et al., 2022] (Fig. 9, Methods Sec. 4.1). We trained this CMM via backpropagation through time (BPTT) to carry out two distinct cognitive paradigms: the visual-change-detection task and an evidence-accumulation task, achieving accuracies similar to those reported for mice (Fig. 6F and Fig. S2D). Crucially, the CMM reproduced the main sequence-level signature observed in V1: sparse activity with peak-latency orders that changed across task outcome and image context (Fig. 3; evidence-accumulation results in Fig. S1).

**Figure 3:**
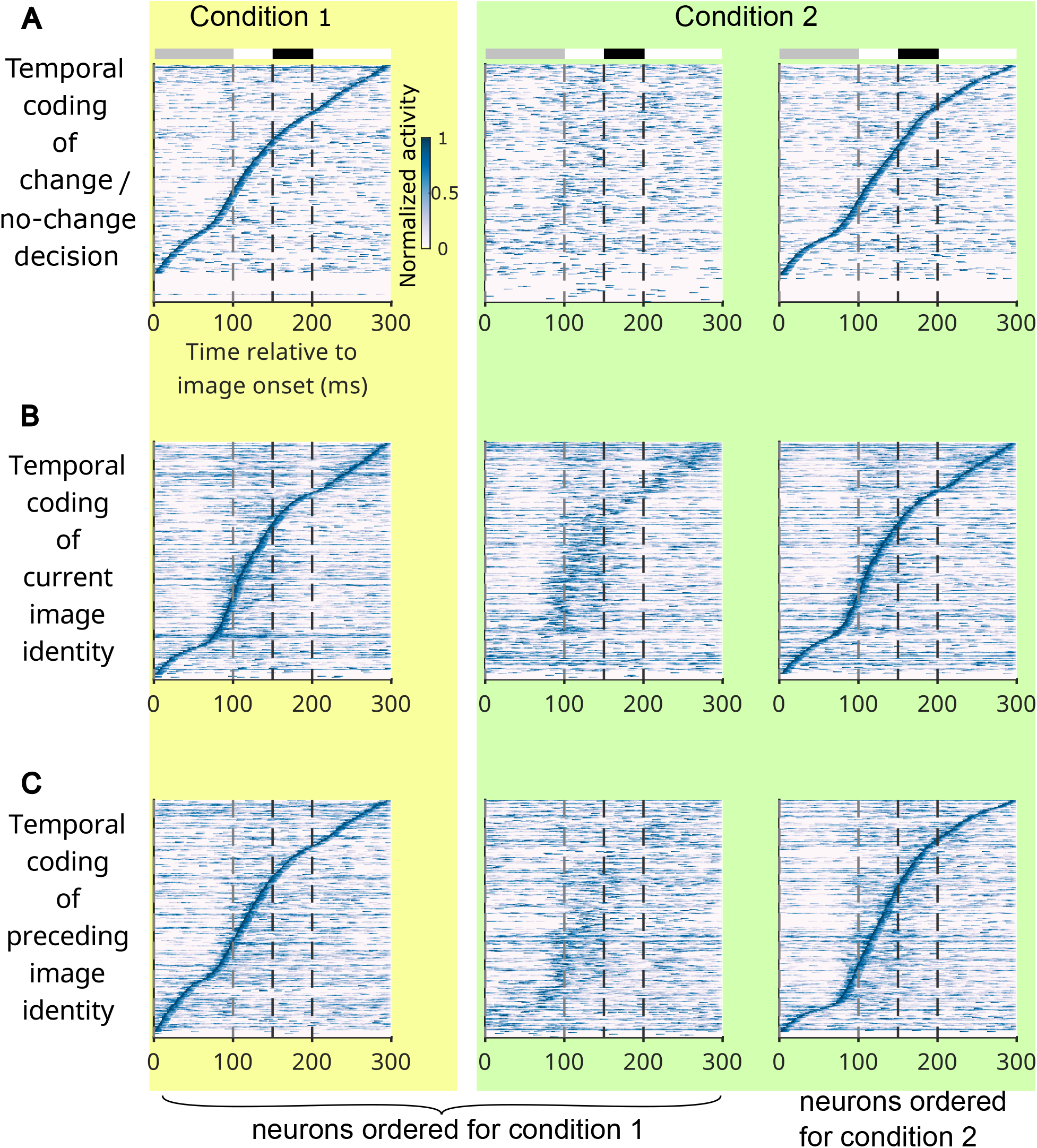
Condition-dependent spike-order structure in the trained CMM during the visual-change-detection task. **(A-C)** Same as in Fig. 2A-C but for neuronal activity in the CMM, which had been trained for this task. Gray and black bars at the top indicate the image presentation and response windows, respectively. As in the mouse V1 recordings in Fig. 2, neural activity is sparse and exhibits a clear sequential organization with high temporal resolution. The blurred sequences in middle panels, where neurons are ordered for condition 1, indicate that the order of peak activity of neurons is quite different for the two conditions.

We then compared the CMM with two matched recurrent controls: a randomly connected recurrent spiking neural network (RSNN) and a rate-based RNN (Methods Sec. 4.7). These controls learned the tasks, but their peak-latency orders were more similar across conditions than those of the CMM and mouse V1 (Fig. 4A,B; quantified in Sec. 2.5). This comparison does not rule out all possible RNN or SNN architectures; it shows that generic recurrence and spiking, under the tested control architectures and training protocol, were insufficient to match the cortical spike-order signature.

**Figure 4:**
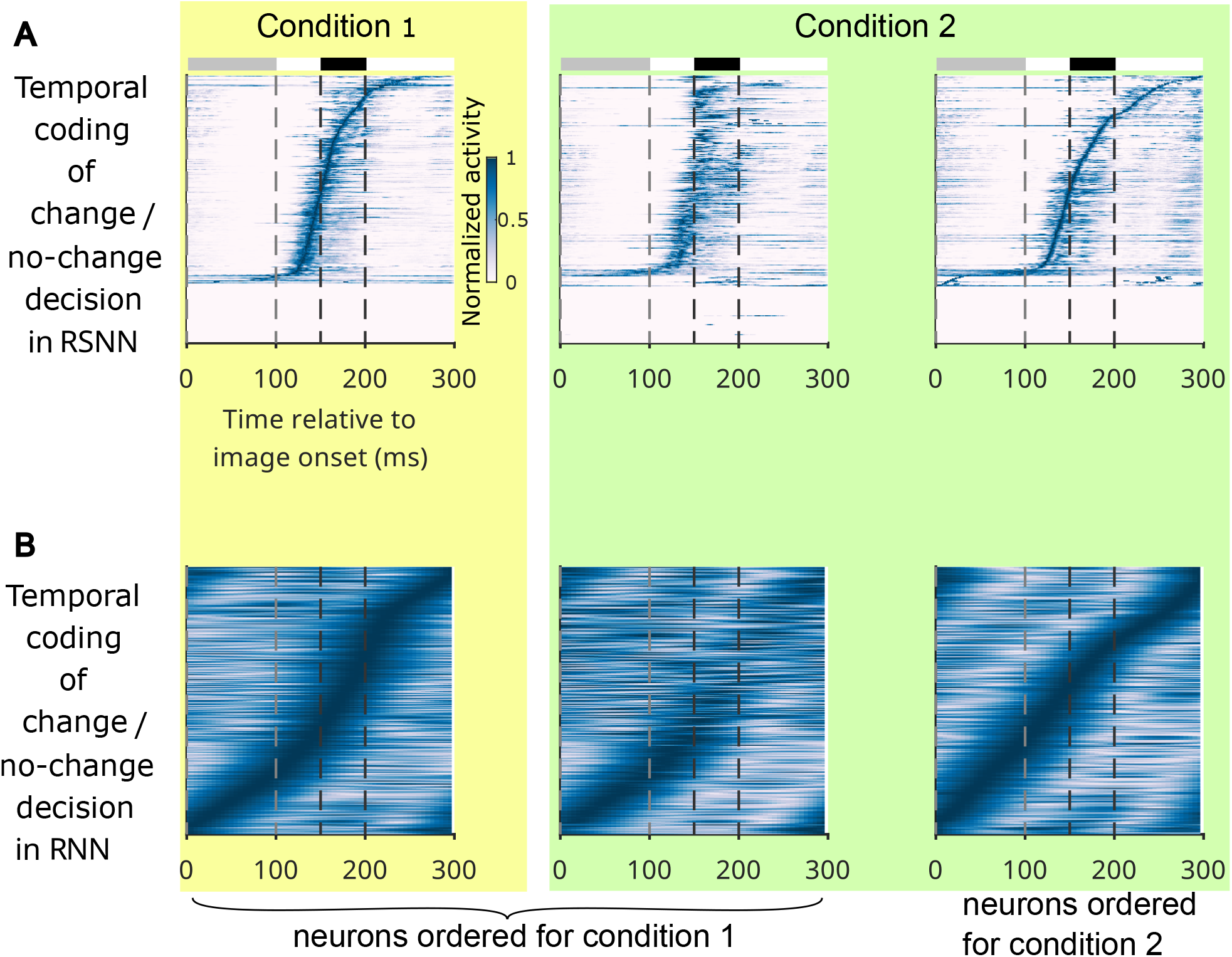
Tested recurrent controls show less condition-dependent spike-order structure. **(A)** Same as in Fig. 3A but for a randomly connected recurrent spiking network (RSNN) matched to the CMM in neuron and synapse counts. Gray and black bars at the top indicates the image presentation and response windows, respectively. **(B)** Same as in **(A)** but for a rate-based recurrent neural network (RNN) matched in size. In contrast to Fig. 2 and Fig. 3, the middle and right panels are more similar, indicating weaker condition dependence of peak-latency order in these tested controls.

### 2.3 Trial-to-trial stability of sparse temporal ordering

A critical challenge for spike-based computation is maintaining reliable outputs despite substantial trial-to-trial variability. The awake neocortex exhibits significant variability in precise spike timing—with individual neurons often shifting their firing across identical trials. This high variability is faithfully replicated by the empirical, data-driven noise model of the CMM, which yields a coefficient of variation closely matching the *in vivo* distribution (Fig. 5B, E, Methods Sec. 4.1.3).

**Figure 5:**
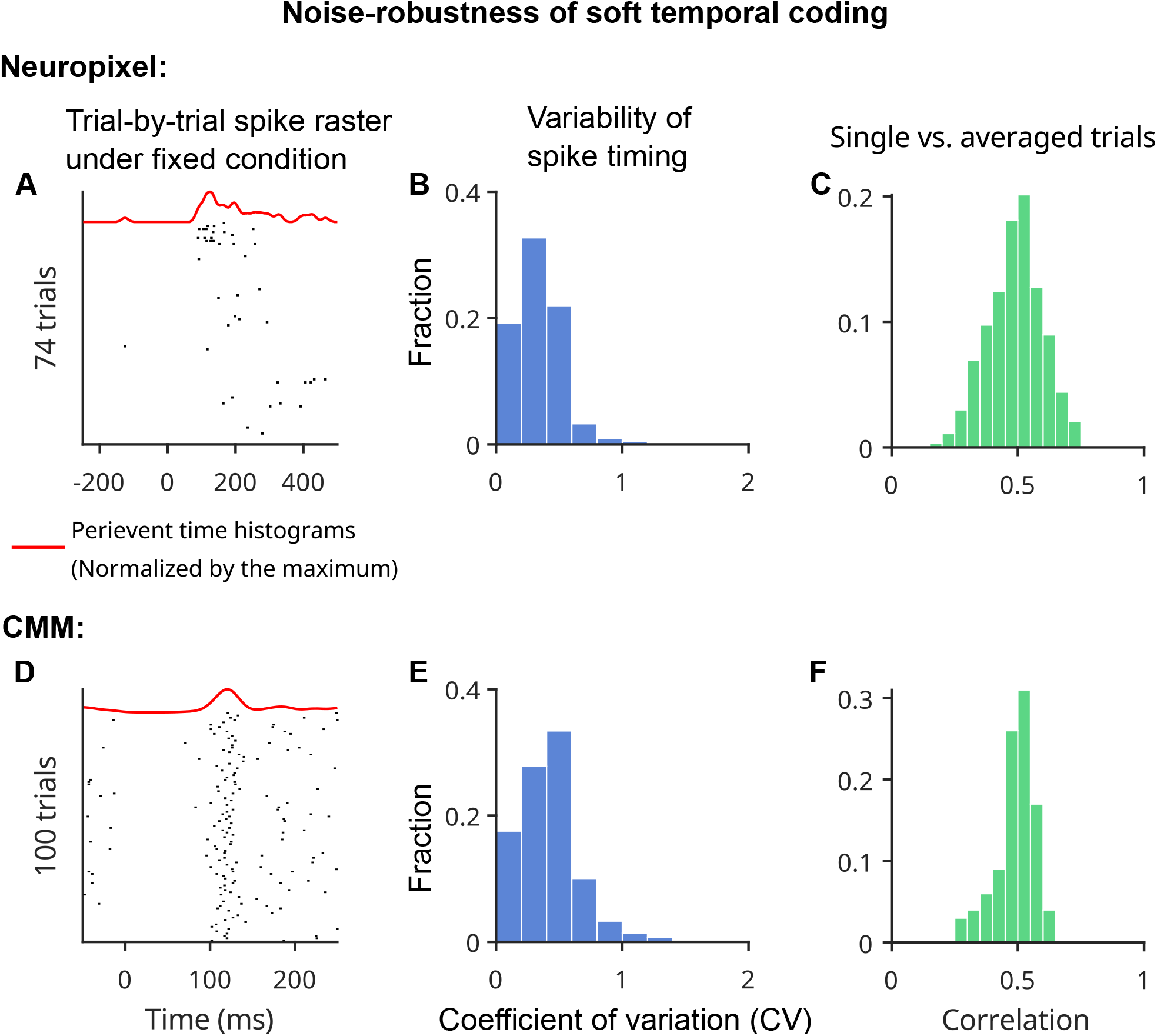
Single-trial variability and stability of soft temporal ordering. **(A)** Trial-by-trial spike raster under fixed condition for a randomly selected Neuropixel recorded neuron in mouse V1 across 74 trials. Each trial has the same preceding and current images. The red curve represents the perievent time histograms normalized by its maximum, smoothed with a 50-ms Gaussian kernel. **(B)** Variability of spike timing in mouse V1 across 74 trials. To quantify trial-to-trial timing jitter, we calculated the coefficient of variation (CV)—defined here as the variability of a neuron’s specific firing time across different trials with identical stimuli. The CV was calculated for the fraction of the 168 recorded neurons with more than two spikes. **(C)** Histogram of Spearman rank correlations between single-trial firing order and trial-averaged firing order. For neurons with multiple spikes in a trial, rank position was determined by mean spike time; non-firing neurons were omitted for that trial. The positive shift (*t*-test, *p* < 10^−10^) indicates reproducible temporal ordering despite timing variability. **(D-F)** Same as **(A-C)**, but for the CMM evaluated across 100 trials.

**Figure 6:**
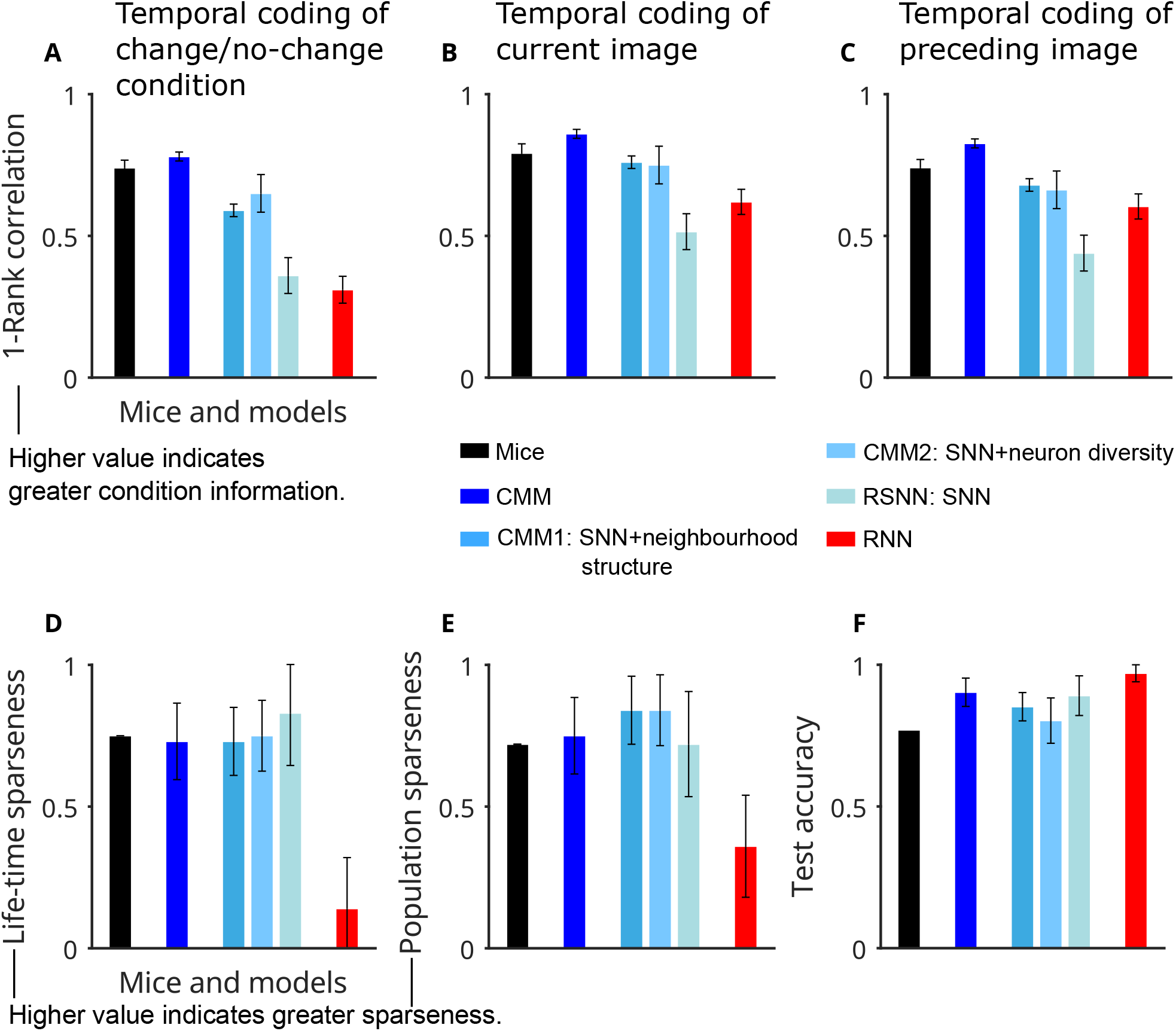
Firing-order condition dependence, sparseness, and task performance across mouse V1 and model variants in the visual-change-detection task. **(A)** 1 - Spearman rank correlation between change and no-change conditions. Larger values indicate greater condition-specific information in firing order. **(B-C)** Same as in **(A)** but for two current-image identities and two preceding-image identities, respectively. **(D)** Comparison of lifetime sparseness among models and mouse V1. Larger values indicate sparser single-neuron responses. **(E)** Same as in **(D)** but for population sparseness. **(F)** Comparison of test accuracy among models and mouse V1. The mouse performance was estimated in Fig. 1I of [Garrett et al., 2020]. **(A-F)** share the same color scheme. Error bars are standard deviations calculated among 10 mice and 10 independently trained model instances.

We next asked whether the condition-dependent sequence structure is compatible with this variability. Both the biological recordings and the CMM are consistent with a “soft” temporal code: the relevant variable is not the millisecond timing of isolated spikes, but the relative order of short spike bouts or mean spike times within a trial (Fig. 5A,D). This bout-based firing might provide a natural buffer against timing jitter.

Furthermore, we confirmed that this sequence organization is not merely an artifact of trial-averaging. In both the biological recordings and the CMM across both tasks, the rank order of spikes in single trials remains strongly and significantly correlated with the average rank order (*t*-test against zero correlation, *p* < 10^−10^) (Fig. 5C, F). These results indicate that sparse temporal ordering can remain reproducible despite substantial spike-timing variability.

### 2.4 Early-informer neurons perturb condition-dependent trajectory separation

To visualize how the trained CMM organizes its high-dimensional activity into these condition-dependent pathways, we embedded exponentially filtered population activity into a low-dimensional state space using PCA followed by UMAP (Methods Sec. 4.9). We observed a trajectory-organization motif across both the visual-change-detection (Fig. 7) and evidence-accumulation tasks (Fig. 8) and make a prediction: whether the network is performing rapid visual comparison or integrating abstract information over time, the underlying dynamical system solves it the same way by steering high-dimensional activity through low-dimensional, task-specific bifurcations. Because this 2D embedding captures only part of the full variance, we use it as a visualization of relative trajectory organization rather than as proof of exact geometric distances in the full state space.

**Figure 7:**
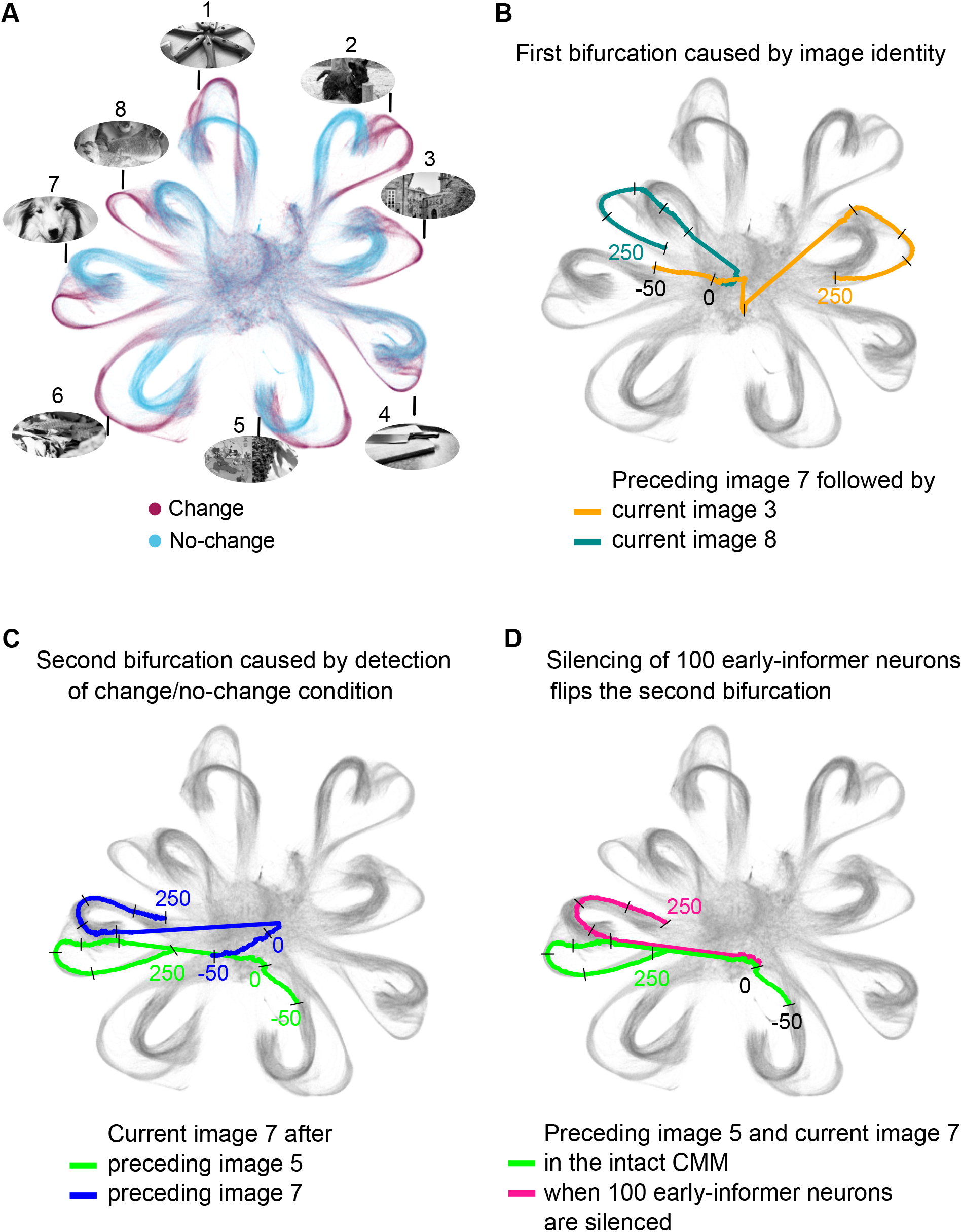
Low-dimensional visualization of condition-dependent CMM trajectories in the visual-change-detection task. Each dot represents network activity at a specific millisecond within a 300-ms fragment of the computation of the CMM on a sequence of natural images. Short black bars mark 50-ms segments of network trajectories. **(A)** Network trajectories of the cortical microcircuit model during the presentation of 8 (out of the 100) test images that had not been presented during training. Two colors indicate whether the current image was identical (blue) or different (purple) from the preceding image. The embedding separates trajectories by image identity and task condition. **(B)** The first bifurcation occurs according to the identity of the current image, highlighted here for the case where the trajectory starts in both cases from the same state, which largely results from the identity of the preceding image. This first bifurcation occurs within the first 50 ms after image onset. **(C)** The second bifurcation does not depend on the identity of the current image, but on whether it is the same or different from the preceding image. The second bifurcation is more difficult to visualize since the trajectory arrives from two different regions that are characteristic of the identity of the preceding image. **(D)** Model-level perturbation of the condition-dependent trajectory. For image 7 after image 5, the intact model trajectory is compared with the trajectory after silencing 100 high-MI early-informer neurons. The displayed altered trajectory is a trial in which the model made an incorrect decision. A corresponding no-change-to-change example is shown in Fig. S4. Note that the displayed altered trajectories represent trials in which the network made incorrect decisions.

**Figure 8:**
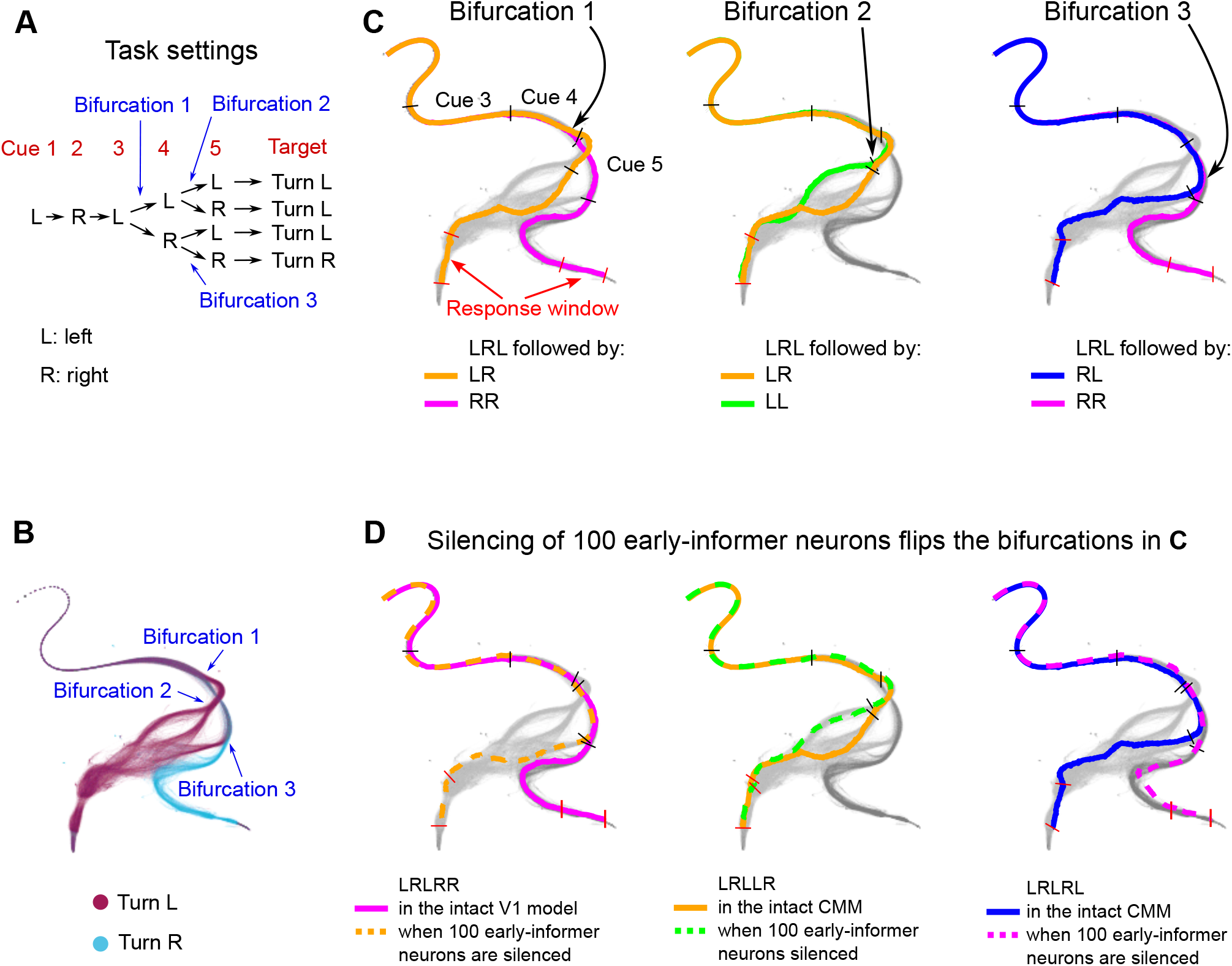
Low-dimensional visualization of condition-dependent CMM trajectories in the evidence-accumulation task. Each dot represents network activity at a particular millisecond of a 530 ms (the first 70 ms was excluded for better visualization) fragment of the computation of the CMM in the evidence-accumulation task. Short black bars mark cues 3-5 on the network trajectories; the red bars denote the response window. **(A)** The task setting. The sequence for the first three cues is consistently left-right-left, while the last two cues can be either left or right. The setting introduces three bifurcations. **(B)** Network trajectories of the CMM during the presentation of five cues. The color differentiation indicates whether the target is to turn left or right. It is evident that the network state trajectories bifurcate three times, depending on the sequence of the fourth and fifth cues. **(C)** The first bifurcation corresponds to the fourth cue, typically occurring 30 ms after its onset. The second bifurcation takes place in response to the fifth cue, provided that the fourth cue is left; however, these two branches eventually converge as they share the same targets. The third bifurcation responds to the fifth cue when the fourth cue is right. **(D)** The bifurcations in **(B)** can be flipped by a small set of early-informer neurons. In each sub-panel, there are two trajectories that have the same fourth and fifth cues in both the intact model and when 100 neurons with the highest mutual information with the current condition of accumulated cues in a temporal segment are silenced. For enhanced visualization, the silenced neurons in the panel from left to right are determined based on the mutual information with the current condition of accumulated cues computed during the fourth, fifth, and fifth cues, respectively. These cues align with the bifurcation pivot points. Note that the displayed altered trajectories represent trials in which the network made incorrect decisions.

**Figure 9:**
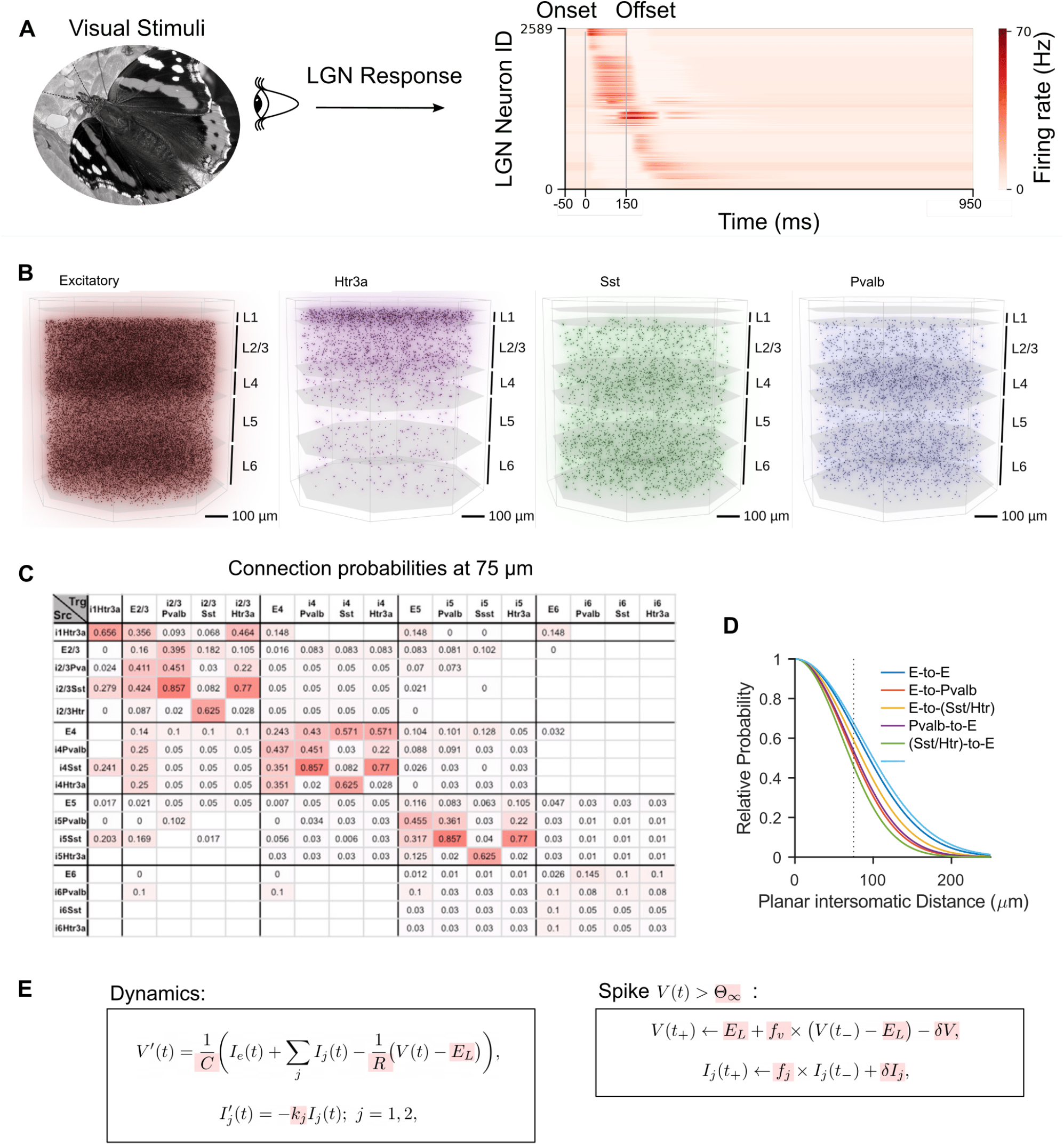
Structure and components of the data-based CMM, which models a column of V1 according to the data of [Billeh et al., 2020]. **(A)** We used the LGN model from [Billeh et al., 2020] to transform visual stimuli into the outputs of 2,589 filters that model firing rates of LGN neurons and are connected to neurons in the cortical microcircuit model according to data-based rules. These outputs provide input currents to V1 neurons using data-based rules. **(B)** Side view of the 3D architecture of the model, which consists of 51,978 neurons with diverse types from 1 excitatory and 3 inhibitory neuron classes (Htr3a, Sst, Pvalb) in a column with 800 µm diameter. These neurons are distributed over 5 laminar sheets (L1 to L6, where L2 and L3 are lumped together). **(C)** The data-based base connection probabilities of [Billeh et al., 2020] depend on the cell class to which the presynaptic (row labels) and postsynaptic neuron (column labels) belongs. White grid cells denote unknown values. **(D)** The base connection probability from **(C)** is multiplied according to [Billeh et al., 2020]) for any given pair of neurons by an exponentially decaying factor that depends on the lateral distance between them. **(E)** Main equations defining the GLIF_3_ neuron models of [Billeh et al., 2020] with 3 internal variables. Assignments to their parameters (highlighted in red) define the 111 neuron models of the networks, based on experimental data from the Allen Brain Atlas [Allen Institute, 2018].

In the visual-change-detection task, trajectories in the embedding separate first according to current image identity and then according to change/no-change condition (Fig. 7B,C). A related sequence of condition-dependent trajectory separations is visible in the evidence-accumulation task (Fig. 8). We refer to these separations descriptively as pathway-like trajectory branches rather than as proof of mathematical bifurcations in the dynamical-systems sense.

Within the trained CMM, we identified neurons with high mutual information between their early activity and the network decision and refer to them operationally as “early-informer” neurons (Methods Sec. 4.8). These neurons were concentrated primarily in layers 2/3 and 4 in the visualization of MI maps (Fig. S3A). Silencing 100 high-MI neurons perturbed the visualized trajectory and caused incorrect decisions in representative trials across both tasks (Fig. 7D, Fig. 8D).

This outsized impact of a small number of neurons might reflect a principle of functional segregation. In the CMM, silencing just 200 neurons reduces performance to chance levels—an effect that requires silencing over 6,000 neurons in a rate-based RNN (Fig. S3B). Notably, generic RSNNs share this extreme sensitivity to small subsets of high-information neurons. However, because standard RSNNs operate using more rigid sequential pipelines, these neurons act more as critical points of failure. This yields a testable prediction for future perturbation experiments: neurons that carry early condition information should disproportionately influence subsequent spike order, trajectory separation, and decisions, although the present results do not establish that homologous biological neurons act as switches *in vivo*.

### 2.5 Architectural constraints associated with flexible temporal coding

The full CMM combines several biological constraints, including neuronal diversity and distance-dependent local connectivity. To determine which of these constraints were associated with the condition-dependent spike-order signatures, we compared the full CMM with matched spiking control models (Table 1). CMM1 preserved the V1-like connectivity pattern but reduced neuronal diversity to one excitatory and one inhibitory type; CMM2 preserved neuronal diversity but randomized recurrent connectivity; and the RSNN lacked both V1-like connectivity and full neuronal diversity. The rate-based RNN was retained as an additional reference model, but was not included in the factorial statistical analysis because it was not part of the same spiking-network design.

**Table 1:** Control model configurations.

| Model Name | Neuron Model | Connectivity<sup>a</sup> | Neuron Diversity<sup>b</sup> | Input via LGN |
| --- | --- | --- | --- | --- |
| CMM | GLIF<sub>3</sub> | V1-like | 111 types | Yes |
| CMM1 | GLIF<sub>3</sub> | V1-like | 1E, 1I types | Yes |
| CMM2 | GLIF<sub>3</sub> | Random | 111 types | Yes |
| RSNN | GLIF<sub>3</sub> | Random | 1E, 1I types | Yes |
| RNN | rate neuron | Random | Homogeneous | Yes |

<sup>a</sup> V1-like: Biologically constrained connectivity of the CMM. Random: Connections randomly assigned while preserving overall synaptic number. <sup>b</sup> 111 types: Full diversity of 111 neuron types. 1E, 1I types: All excitatory neurons are identical (L2/3 E-type, ID:487661754), all inhibitory neurons are identical (L2/3 PV-type, ID:484635029). Homogeneous: All neurons are identical units.

We quantified temporal-code flexibility as rank-order dissimilarity, *D* = 1 − *ρ*, where *ρ* is the Spearman rank correlation between the peak-latency orders of two conditions. Larger values of *D* indicate stronger condition dependence of temporal ordering. The four spiking models formed a balanced 2 × 2 factorial design with Connectivity (V1-like versus random) and Diversity (full neuronal diversity versus reduced excitatory/inhibitory types) as orthogonal factors, with *N* = 10 independently trained model instances per cell.

Across all three temporal-order comparisons, both V1-like connectivity and neuronal diversity significantly increased rank-order dissimilarity after Holm correction (Table S1). This pattern was observed for task outcome, current-image identity, and preceding-image identity, indicating that both architectural constraints contribute robustly to condition-dependent temporal organization. The full CMM showed the strongest temporal-order flexibility across the three comparisons within this spiking ablation family, whereas CMM1 and CMM2 showed intermediate values and the RSNN showed the most rigid temporal ordering (Fig. 6A–C). Thus, removing either neuronal diversity or V1-like connectivity reduced the cortical-like temporal-order signature, although neither ablation abolished it completely.

The Connectivity × Diversity interactions did not survive correction for any temporal-order metric (Table S1). We therefore do not interpret these data as evidence for a strict synergistic or jointly necessary mechanism. Instead, the results support a complementary-constraints model: neuronal diversity and spatially structured local connectivity make separable, largely additive contributions to flexible spike-order coding. In this view, each constraint improves temporal-code flexibility through a partially independent architectural route, and their combination in the full CMM yields the most consistently flexible spiking model tested here.

In contrast, the same factorial analysis did not identify either factor as a reliable determinant of sparseness. Neither lifetime sparseness nor population sparseness was significantly explained by Connectivity, Diversity, or their interaction after correction (Table S1). This dissociation indicates that the architectural constraints associated with flexible temporal ordering are not the primary determinants of sparse activity in these trained networks.

Together, these results suggest a division of labor between activity constraints and architectural constraints. Spiking dynamics and activity regularization help establish a sparse firing regime, whereas neuronal diversity and spatially structured local connectivity promote condition-dependent temporal organization. Sparse firing is therefore compatible with spike-order coding, but is not sufficient to reproduce the cortical temporal-order signature: the RSNN approached the CMM in sparseness while showing weaker condition-dependent temporal ordering, whereas the rate-based RNN remained in a denser and more rigid activity regime (Fig. 6D,E).

## 3 Discussion

The central message of this study is that sparse cortical computation need not be temporally rigid. Across large-scale recordings from mouse cortex during visual change detection and evidence accumulation [Garrett et al., 2020, Arkhipov et al., 2025, Koay et al., 2022], task-relevant information was reflected not only in which neurons fired, but in how their peak activity was ordered over time. This order was flexible: it changed with task outcome, current stimulus identity, and preceding stimulus identity (Fig. 2). Thus, the code observed here lies between two familiar extremes. It is richer than a static sparse rate code, because the same sparse population can be reorganized into different temporal orders. But it is also more tolerant than a strict millisecond spike-timing code, because the relevant structure appears at the level of short spike bouts and relative firing order rather than exact spike times (Fig. 5). In this sense, the data support a soft spike-order code: a representational format in which sparse spikes remain reliable enough to support computation, yet flexible enough to route different task conditions through different temporal pathways.

This interpretation connects with, but also refines, earlier views of temporal coding in spiking networks. Classical precise-timing hypotheses emphasized the computational power of exact spike times [Thorpe, 1990, Maass, 1994]. The present results do not argue against temporal coding; rather, they suggest that cortex may use a more robust form of it. Instead, what matters is the relative order of population events, a coding scheme that can survive biological noise and trial-to-trial variability. This distinction is important for both neuroscience and artificial SNN design.

In order to understand how this flexible, condition-dependent computation can be ported into artificial SNNs, we employed as a research tool a detailed computer model for a generic cortical microcircuit (CMM) [Mountcastle, 1998, Douglas and Martin, 2004], specifically the model for a cortical column in area V1 of [Billeh et al., 2020, Chen et al., 2022], see Fig. 9. After task optimization, the CMM reproduced the key combination seen in the recordings: sparse firing together with condition-dependent temporal order (Fig. 3). Compared with the matched randomly connected RSNN and rate-RNN controls, the CMM occupied a more cortical-like regime, in which computation was neither dense and rate-like nor locked into a single rigid sequence (Figs. 4 and 6). Further ablation results demonstrate that neuronal diversity and distance-dependent local connectivity both increased rank-order dissimilarity across task outcome, current image, and preceding image (Fig. 6 and S2). A useful interpretation is that cortical architecture supplies multiple complementary inductive biases. Heterogeneous neurons create diverse intrinsic timescales and response nonlinearities; local connectivity constrains how activity can propagate through space and cell classes. Together, these features make it harder for training to collapse the network into a single stereotyped temporal pipeline, and easier for different task contexts to recruit different spike-order patterns. Importantly, these architectural ingredients are not exotic additions to artificial SNNs: neuronal heterogeneity and distance-dependent connectivity can be implemented as modest extensions of standard randomly connected RSNNs, and are compatible in principle with current neuromorphic platforms such as Intel’s Loihi 2 [Shrestha et al., 2024, Rao et al., 2022].

This dissociation between sparseness and temporal flexibility is one of the most important conceptual outcomes of the study. A network can be sparse without being cortical-like. The RSNN approached the CMM in some measures of sparseness, yet showed more rigid temporal ordering; the rate-RNN remained denser and less spike-order-like (Fig. 6). Thus, sparseness is better viewed as a necessary operating regime than as a complete computational mechanism. Activity regularization and spiking dynamics can make a network economical, but architectural structure determines whether this economy is used as a flexible routing substrate or merely as a compressed sequential pipeline. For neuromorphic computing, this distinction matters: reducing spikes saves energy, but organizing the remaining spikes into condition-dependent pathways is what may provide computational flexibility.

The low-dimensional trajectory analyses offer a second view of the same principle. In both tasks, population activity in the trained CMM separated into condition-dependent trajectory branches in UMAP embeddings (Figs. 7 and 8). The low-dimensional trajectory analyses reveal how the temporal code can be read dynamically: the same task variables that reorder sparse spike sequences also separate population trajectories. Spike-order flexibility and trajectory branching therefore appear to be two descriptions of a common routing phenomenon, one observed in the timing of sparse events and the other in the geometry of evolving population states.

The early-informer analysis further suggests how such routing could become causally controllable. In the CMM, neurons whose early activity carried high mutual information about the later decision had a disproportionate influence on trajectories and behavior when silenced (Figs. 7, 8, S3, and S4). We use the term early-informer neurons operationally, not as a claim that an identical biological cell class has already been identified. The model-derived prediction is sharper: within sparse cortical computations, small subsets of neurons that fire early and carry condition information should act as control points for later trajectory selection. This prediction is consistent with experimental demonstrations that cortical activity and behavior can be sensitive to small neuronal subsets [Vinje and Gallant, 2000, Houweling and Brecht, 2008, Doron et al., 2014, Marshel et al., 2019, Dalgleish et al., 2020, Doron et al., 2020], but it remains to be tested directly for the spike-order and trajectory-routing signatures proposed here.

Interestingly, these lessons from cortical spiking computation have striking parallels with algorithmic and architectural strategies that have recently been proposed in artificial intelligence: Pathway computing [Dean, 2021, Gesmundo and Dean, 2024] and modern mixtures of experts (MoE) [Fedus et al., 2022, Muennighoff et al., 2024, Abnar et al., 2025]. These methods are used in large LLMs to extract task-relevant information from billions of parameters without activating the entire network. The insights derived from mouse V1 circuits suggest a biologically motivated route for implementing related routing principles in smaller, energy-efficient SNNs for edge settings.

Several limitations define the scope of this interpretation. Peak-latency rank order is an operational measure of temporal organization and does not by itself prove anatomical pathway switching in the biological circuit. UMAP visualizations reveal low-dimensional organization but cannot determine the full geometry of the neural state space. The CMM simplifies many biological factors, including plasticity, long-range inputs, neuromodulation, behavioral state, and developmental constraints. The controls tested here also do not exhaust the space of possible SNNs, RNNs, or modern machine-learning architectures. Future experiments could test the strongest prediction more directly by perturbing candidate early-informer neurons during specific task epochs and asking whether this selectively changes the subsequent spike order, trajectory branch, and behavioral decision.

Taken together, the results point to a cortical design principle for efficient computation: sparseness is not merely a way to reduce activity, but a substrate for condition-dependent temporal routing. Sparseness provides the economy; cellular diversity and local connectivity provide the structure that makes sparse activity flexible. This view offers a compact lesson for brain-inspired AI: copying every biological detail is unnecessary, but copying the right architectural constraints may allow artificial SNNs to inherit a central cortical advantage–the ability to compute through sparse, flexible, condition-dependent temporal pathways.

## 4 Methods

### 4.1 Cortical microcircuit model

In contrast to standard recurrent neural networks (RNNs) with homogeneous units and random connectivity, the CMM provides a more faithful representation of a cortical column, receiving inputs from a simple LGN model (Fig. 9A, Sec. 4.2 of Methods).

The model comprises 51,978 spiking neurons distributed across a 3D cylindrical volume, replicating the laminar structure of the neocortex. This model is distinguished by its incorporation of extensive biological constraints drawn from experimental data (Fig. 9). Critically, these are not uniform units; the CMM incorporates 17 different neuron types (listed in each row and column of Fig. 9B). These neuron types are further split into 111 different variations based on response profiles of individual neurons [Allen Institute, 2018] of the neocortex, each parameterized by a generalized leaky integrate-and-fire (GLIF_3_) model [Teeter et al., 2018] that was fitted to electrophysiology recordings (Fig. 9E). Furthermore, synaptic connectivity is not random but is governed by data-driven probabilistic rules that depend on both the pre- and post-synaptic neuron types and their 3D spatial distance (Fig. 9C, D). This structure results in a network dominated by local connections, a hallmark of cortical organization.

#### 4.1.1 Neuron models

As in [Chen et al., 2022], we focused on the “core” part of the point-neuron version of the realistic CMM introduced by [Billeh et al., 2020]. It contains 51,978 neurons in a 400-*µ*m-radius column. We also tested a smaller column radius and found that even in a column containing 1,000 neurons our results are still valid. To make it more gradient-friendly, we replaced the hard reset of membrane potential after a spike emerges with the reduction of membrane potential (soft reset) *z*_*j*_(*t*) (*v*^th^ − *E*_L_), where *z*_*j*_(*t*) = 1 when neuron *j* fires at time *t* and *z*_*j*_(*t*) = 0 otherwise. *v*^th^ is the firing threshold of membrane potential and *E*_*L*_ the resting membrane potential. Soft reset causes no significant change in the neural response [Chen et al., 2022]. We simulated each trial for 600 ms for both visual-change-detection and evidence-accumulation tasks. The dynamics of the modified GLIF_3_ model was defined as

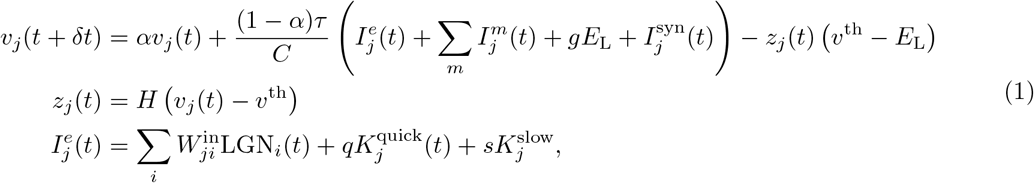

where *C* represents the neuron capacitance, *I*^*e*^ the external current, *I*^syn^ the synaptic current, *g* the membrane conductance, and *v*^th^ the spiking threshold. 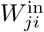 is the synaptic weight from LGN neuron *i* to V1 neuron *j*. The scales of the quick noise 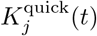 and the slow noise 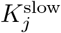 to neuron *j* are *q* = 2 and *s* = 2, respectively, unless otherwise stated. *K*_*j*_ was randomly drawn from the empirical noise distribution which will be elaborated on later. The decay factor *α* is given by *e*^−*δt/τ*^, where *τ* is the membrane time constant. *δt* denotes the discrete-time step size, which is set to 1 ms in our simulations. *H* denotes the Heaviside step function. To introduce a simple model of neuronal refractoriness, we further assumed that *z*_*j*_(*t*) is fixed to 0 after each spike of neuron *j* for a short refractory period depending on the neuron type. The after-spike current *I*^*m*^(*t*) was modeled as

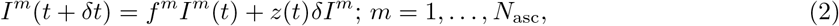

where the multiplicative constant *f* ^*m*^ = exp (−*k*^*m*^δ*t*) and an additive constant, *δI*^*m*^. In our study, *m* = 1 or 2. Neuron parameters have been fitted to neurophysiological data from 111 selected neurons according to the cell database of the Allen Brain Atlas [Allen Institute, 2018], see [Teeter et al., 2018, Billeh et al., 2020], including neuron capacity *C*, conductance *g*, resting potential *E*_L_, firing threshold *v*^th^, the length of the refractory period, as well as amplitudes *δI*^*m*^ and decay time constants *k*^*m*^ of two types of after-spike currents, *m* = 1, 2.

#### 4.1.2 Synaptic inputs

The CMM utilizes experimental data to specify the connection probability between neurons. The base connection probability for any pair of neurons from the 17 cell classes is provided in [Billeh et al., 2020] in a table (shown in Fig. 9B), where white cells denote unknown values. The values in this table are derived from measured frequencies of synaptic connections for neurons at maximal 75 µm horizontal inter-somatic distance. The base connection probability was then scaled by an exponentially decaying factor based on the horizontal distance between the somata of the two neurons (Fig. 9C), also derived from experimental data [Billeh et al., 2020]. The synaptic delay was spread in [1, 4] ms, as extracted from Fig. 4E of [Billeh et al., 2020] and rounded to the nearest integer as the integration step is 1 ms.

The postsynaptic current of neuron *j* was defined by the following dynamics [Billeh et al., 2020]:

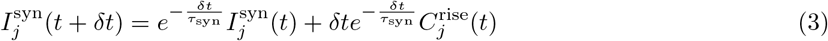

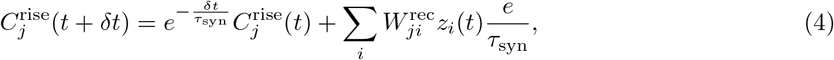

where *τ*_syn_ is the synaptic time constant, 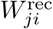is the recurrent input connection weight from neuron *i* to *j*, and *z*_*i*_ is the spike of presynaptic neuron *i*. The *τ*_syn_ constants depend on neuron types of pre- and postsynaptic neurons [Billeh et al., 2020].

#### 4.1.3 Data-driven noise model

We used a noise model that was introduced in our previous study [Chen et al., 2022]. The model was based on an empirical noise distribution that was obtained from experimental data of mice responses to 2,800 nature images [Stringer et al., 2019]. The noise currents 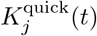 and 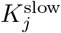 in Eq. 1 were drawn independently for all neurons from this distribution. The quick noise 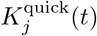 was drawn every 1 ms while the slow noise 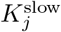 was drawn once every 600 ms. The empirical noise distribution was derived from the variability (additive noise) collected from the mouse V1 data. A detailed mathematical analysis of this method is available in the methods and supplementary materials of [Stringer et al., 2019].

A critical feature of the CMM is its replication of the strong, pervasive noise found in mouse V1. This realistic noise injects significant trial-to-trial variability into the network’s spiking activity, often shifting spike timing by several dozens of milliseconds and altering the precise number of spikes a neuron emits per trial (Fig. 5A, D). We quantified this variability, finding a coefficient of variation (CV) of 0.32 ± 0.18 (mean±std) in Neuropixel recordings and 0.42 ± 0.24 in the CMM (Fig. 5B, E).

#### 4.1.4 Projection neurons

Unlike the conventional method of using linear readouts from all neurons, which can obscure the micro-circuit’s computational contributions (only training the linear readout could get already high accuracy) [Maass et al., 2002, Maass and Markram, 2004, Chen et al., 2022], we randomly selected a single pyramidal neuron in layer 5 of the CMM to serve as projection neurons (Fig. S3A), responsible for reporting network decisions through their firing patterns [Chen et al., 2022]. We also tested 10, 30, 50 and 100 projection neurons and found that the number of projection neurons does not affect the results; all reported results use a single projection neuron. This approach aligns with the biological role of pyramidal neurons in transmitting computational outcomes to other brain areas [Harris and Shepherd, 2015, Marshel et al., 2019], triggering behavioral responses.

### 4.2 Training task and injecting visual input into the model

#### LGN model

The visual stimuli were processed by a qualitative retina and LGN model, following [Billeh et al., 2020]. Their full LGN model consists of 17,400 spatiotemporal filters that simulate the responses of LGN neurons in mice to visual stimuli [Durand et al., 2016]. Each filter generates a positive output, which represents the firing rates of a corresponding LGN neuron. We used only a subset of 2,589 of these LGN filters that provide inputs from a smaller part of the visual field to the core part of the CMM, on which we are focusing in this study. The input images were first converted to grayscale and scaled to fit in the interval [−*Int, Int*], where *Int* > 0. The output of the LGN model was then used as an external current input in the CMM as follows:

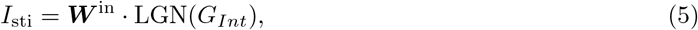

where *G*_*Int*_ represents images scaled into [−*Int, Int*] for *Int* = 2.

#### Visual-change-detection task with natural images

We designed the visual-change-detection task to be as close as possible to corresponding biological experiments while keeping them as simple as possible. In the mouse experiments of [Garrett et al., 2020, Siegle et al., 2021], mice were trained to perform a visual-change-detection task using static natural images presented in a sequence of 250 ms with short phases (500 ms) of gray screens in between. The mice had to report whether the most recently presented image was the same as the previously presented one. To replicate this task while taking into account GPU memory limitations, we presented natural images for 100 ms each with delays between them lasting 200 ms (Fig. 1B). The first image was presented after 50 ms, and all images were selected from a set of 40 randomly chosen images from the ImageNet dataset [Deng et al., 2009]. The model had to report within a 50 ms time window starting 150 ms after image onset (response window) if the image had changed.

We randomly selected a pool of 140 natural images from the ImageNet dataset [Deng et al., 2009] that we used as network inputs. Similar to the biological experiments of [Garrett et al., 2020, Siegle et al., 2021], we used 40 of them for training (familiar images), and 100 of them for testing (novel images).

In the response window, we defined the mean firing rate of readout population as

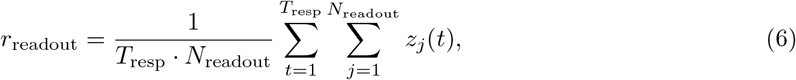

where the sum over *j* is over the *N*_readout_ projection neurons and the sum over *t* is over the time length of response window *T*_resp_ = 50 ms. If *r* > *r*_0_ = 0.01, the model reported a network decision that the image had changed. Otherwise, it reported no change.

#### evidence-accumulation task

A hallmark of cognitive computations in the brain is the capability to go beyond a purely reactive mode: to integrate diverse sensory cues over time, and to wait until the right moment arrives for an action. A large number of experiments in neuroscience analyze neural coding after learning such tasks (see e.g., [Morcos and Harvey, 2016, Engelhard et al., 2019]). We considered the same task that was studied in the experiments of [Morcos and Harvey, 2016, Engelhard et al., 2019]. There a rodent moved along a linear track in a virtual environment, where it encountered several visual cues on the left and right. Later, when it arrived at a T-junction, it had to decide whether to turn left or right. The network should report the direction from which it had previously received the majority of visual cues. To reproduce this task under the limitations of a GPU implementation, we used a shorter duration of 600 ms for each trial (Fig. 1C). The right (left) cue was represented by 50 ms of cue image in which the black dots appear on the right (left) side of the maze. Visual cues were separated by 10 ms, represented by the gray wall of the maze. After a delay of 250 ms, the network had to decide whether more cues had been presented on the left or right, using two readout populations for left and right. The decision was indicated by the more vigorously firing readout pool (left or right) within the response window of 50 ms.

#### Injecting image without LGN

As a control measure, we also trained models in the absence of LGN. Echoing the no-LGN input approach for control model 1 in Ref. [Chen et al., 2022], we fed the image’s pixel values directly into the CMM. This image was resized to approximate the number of active LGN channels. The pixel values were multiplied by a factor of 0.04 to align the firing activity within a suitable range. Without LGN, none changed the qualitative conclusion.

### 4.3 Loss function

The loss function was defined as

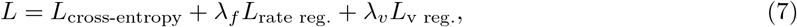

where *L*_cross-entropy_ represents the cross-entropy loss, *λ*_*f*_ and *λ*_*v*_ represent the weights of firing-rate regularization *L*_rate reg._ and voltage regularization *L*_v reg._, respectively. As an example, the cross-entropy loss of visual-change-detection task was given by

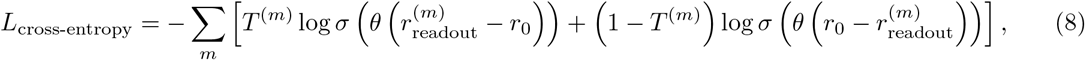

where the sum over *m* is organized into chunks of 50 ms and 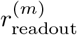 denotes the mean readout population firing rate defined in Eq. 6. Similarly, *T* ^(*m*)^ denotes the target output in time window *m*, being 1 if a change in image identity should be reported and otherwise 0. The baseline firing rate *r*_0_ was 0.01. *σ* represents the sigmoid function. *θ* is a trainable scale (*θ* > 0) of firing rate.

We used regularization terms in the loss function to penalize very high firing rates as well as values of membrane voltages that were not biologically realistic. The default values of their weights were *λ*_f_ = 0.1 and *λ*_v_ = 10^−5^. The rate regularization is defined via the Huber loss [Huber, 1992] between the target firing rates, *y*, calculated from the model in [Billeh et al., 2020], and the firing rates, *r*, sampled the same number of neurons from the network model:

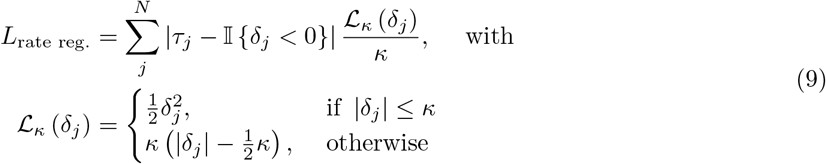

where *j* represents neuron *j, N* the number of neurons, *τ*_*j*_ = *j/N*, *δ* = 0.002, 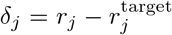. 𝕀 (*ζ*) = 1 when *ζ* is true; otherwise 0.

The voltage regularization is defined through the term

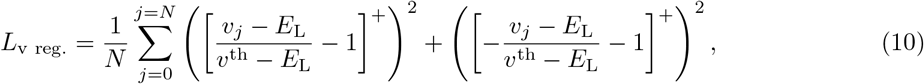

where *N* represents the total number of neurons, *v*_*j*_, the membrane potential of neuron *j*, […]^+^, the rectifier function. *v*^th^ is the firing threshold of membrane potential. *E*_*L*_ the resting membrane potential.

### 4.4 Training and testing

We applied backpropagation through time (BPTT) [Chen et al., 2022] to minimize the loss function. We use BPTT not as a model of biological learning, but as a powerful optimization tool to explore the computational capabilities inherent in the CMM’s brain-like structure. The solutions it discovers reveal the potential of this architecture to support complex, efficient dynamics. The non-existing derivative 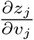 was replaced in simulations by a simple nonlinear function of the membrane potential that is called the pseudo-derivative. Outside of the refractory period, we chose a pseudo-derivative of the form

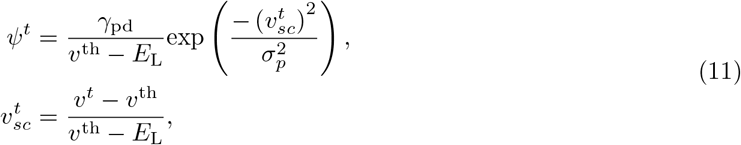

where the dampening factor *γ*_pd_ = 0.5, the Gaussian kernel width *σ*_*p*_ = 0.28. During the refractory period, the pseudo derivative was set to 0.

To demonstrate how sensitive the performance is to the scale of the surrogate derivative, we trained the model with *γ*_pd_ = 0.25 and 0.75 and kept all other hyperparameters the same. When *γ*_pd_ = 0.25, the task performance (testing accuracy) is 0.7; when *γ*_pd_ = 0.75, the testing accuracy is 0.75. Compared with the case of *γ*_pd_ = 0.5 where the task performance is 0.83, other values are worse. This demonstrates that the choice of the derivative’s scale can substantially affect gradient-based learning performance in spiking neural networks [Zenke and Vogels, 2021].

We drew a batch of visual stimuli (64) and calculated the gradient for each synaptic weight to determine whether an increase or decrease of it (but without changing its sign) would reduce the loss function. Weights were then updated by the average gradient across the batch. This method had originally only been applied to neuron networks with differentiable neuron models and was normally referred to as stochastic gradient descent. We used the Adam optimizer with a learning rate of 0.001. Backpropagation was computed over the full trial length without truncation. Unless specified otherwise, initial conditions for spikes were set to zero, and membrane potentials were initialized to the resting potential. The initial configurations for **W**^in^ and **W**^rec^ were adopted from [Billeh et al., 2020]. During the training, we added the sign constraint on the weights of the neural network to keep Dale’s law. Specifically, if an excitatory weight was updated to a negative value, it would be set to 0; vice versa. In every training run, we used a different random seed in order to draw fresh noise samples from the empirical distribution, and to randomly generate/select training samples.

The BPTT training algorithm was implemented in TensorFlow, which is optimized to run efficiently on GPUs, allowing us to take advantage of their parallel computing capabilities. We distributed the visual-change-detection task trials over batches, with each batch containing 64 trials, and performed independent simulations in parallel. Each trial lasted for 600 ms of biological time, and computing gradients for each batch took around 5 s on an NVIDIA A100 GPU. Once all batches had finished (one step), gradients were calculated and averaged to update the weights by BPTT. We define an epoch as 500 iterations/steps. This computation had to be iterated for 22 epochs to make sure the performance was saturated. This took 12 hours of wall clock time on 32 GPUs.

### 4.5 Quantifying discrepancies in sequential patterns

We used Spearman’s rank correlation as an operational measure of similarity between two peak-latency orders. A low correlation between two conditions indicates that neurons change their relative peak-latency order across those conditions. This metric quantifies condition dependence of temporal ordering; it does not by itself identify causal routing or anatomical pathways. For two rank vectors (*R*(*ν*_*i*_), *R*(*µ*_*i*_)), Spearman’s rank correlation coefficient *ρ* is

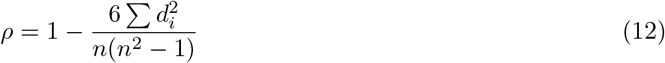

where *d*_*i*_ is the difference between the ranks of corresponding values of *ν*_*i*_ and *µ*_*i*_ (*d*_*i*_ = *R*(*ν*_*i*_) − *R*(*µ*_*i*_)), and *n* is the number of observations.

To ensure that the observed sequential activity was not a sorting artifact, we performed a split-half cross-validation across all 200 trials. The trials were randomly partitioned into two independent halves (fold A and fold B). The neuron sorting order was first established based on peak latency in fold A and subsequently applied to fold B. To evaluate the stability of the sequences, we calculated the Spearman rank correlation between the resulting sequences of the two folds, with statistical significance determined via a *t*-test.

For analyzing single trial rank order, in each trial, we computed a mean spike time for each firing neuron and compared the sequence of neuron firing in that trial with the sequence derived from averaged activity using Spearman’s rank correlation. The averaged activity profile in Figs. 5AD, corresponding to the perievent time histogram (PETH), defines the sequence for this comparison. For single-trial sequence, given that neurons may emit more than one spike in a single trial, we calculated the mean spike time for each firing neuron within each trial. The spike time of non-firing neurons is set as “not a number (nan)” and ignored in the correlation calculation. To confirm that the distribution of rank correlations across trials was significantly different from zero, we employed a *t*-test (*p* < 10^−10^). This analysis underscores the presence of a consistent temporal pattern within individual trials, aligning positively with the averaged activity pattern.

### 4.6 Sparseness

Lifetime sparseness of a neuron is commonly measured according to [Rolls and Tovee, 1995, Vinje and Gallant, 2000, de Vries et al., 2020] by the quotient:

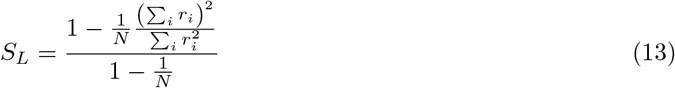

In our experiment, the terms in Eq. 13 are defined as follows. The number of stimulus conditions is *N* = 100, corresponding to 100 unique natural images in test set. A single “trial” consisted of the presentation of one image for a 100 ms duration, followed by a 200 ms inter-trial interval showing a blank gray screen. The neural response, *r*, is quantified as the spike count within a stimulus presentation window set. The term *r*_*i*_ represents the average response of the neuron to the *i*-th image, calculated across all repeated trials for that specific stimulus.

Population sparseness is measured by a similar formula, but the terms are redefined: *N* becomes the total number of neurons (51,978 in the RNNs, CMM and its variants) in the recorded population, and *r*_*i*_ becomes the response of the *i*-th neuron, averaged across all 100 stimulus conditions.

### 4.7 Control Models

To evaluate the specific contributions of CMM features, we employed several control models. This section details the standard Recurrent Neural Network (RNN) used for comparison and other CMM variants. The four primary control models are summarized in Table 1.

#### CMM’s control models

Further control models were derived by selectively modifying the full CMM:

- CMM1: The 111 distinct GLIF_3_ neuron types of the CMM were replaced by a single generic excitatory neuron type (L2/3 excitatory, Allen Brain Atlas ID: 487661754) and a single generic inhibitory neuron type (L2/3 PV, Allen Brain Atlas ID: 484635029).
- CMM2: The CMM’s laminar connectivity was replaced with an equal number of randomly chosen connections, preserving overall connection number but removing spatial structure.

Feedforward connections from the LGN model to V1 were left unchanged. These control models were trained identically to the full CMM, using the same tasks, loss function (including sparsity regularization), and hyperparameters. Specific configurations are detailed in Table 1.

#### Recurrent neural network (RNN) model

The standard rate-based RNN dynamics were defined as:

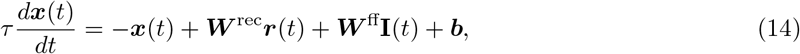

where ***x***(*t*) represents the activation vector of network units at time *t*. The corresponding “firing rate” vector ***r***(*t*) was defined by the element-wise hyperbolic tangent function:

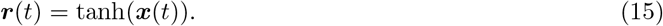

The single-unit timescale *τ* was set to 50 ms, consistent with prior work [Sussillo et al., 2015, Pollock and Jazayeri, 2020]. ***W*** ^rec^ is the recurrent synaptic weight matrix, ***W*** ^ff^ is the feedforward input weight matrix, and ***b*** is the bias vector. For qualitative comparison with CMM activity in Fig. 3A, the absolute value of ***r***(*t*) was used, as ***r***(*t*) ∈ [−1, 1]^*N*^. Weights (***W*** ^rec^, ***W*** ^ff^) and biases (***b***) were initialized from a Gaussian distribution *𝒩* (0, *σ*^2^). For ***W***^rec^ and 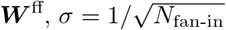 where *N*_fan-in_ is the average number of afferent connections per neuron. For ***b***, *σ* = 0.1. The input **I**(*t*) was preprocessed by the LGN model (Sec. 4.2) unless otherwise specified; using raw stimulus inputs did not yield observable differences in the RNN’s capacity to emulate CMM behaviors. The network’s decision was determined by a linear readout:

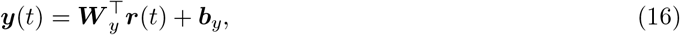

where ***W*** _*y*_ is an *N* × *D*_out_ readout weight matrix (*N* is the number of recurrent neurons, *D*_out_ = 2 for binary decisions), and ***b***_*y*_ is the readout bias. The number of neurons (*N*) and the total number of synaptic connections in the RNN were matched to those in the CMM. The connectivity structure of ***W*** ^rec^ was random (connections drawn uniformly without replacement to match synaptic number) unless stated otherwise.

The RNN loss function was:

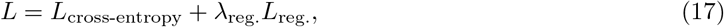

where *L*_cross-entropy_ is defined in Eq. 8 (same for CMM), and *L*_reg._ is an activation regularization term:

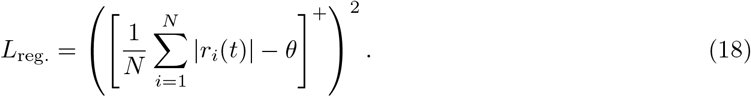

Here, *r*_*i*_(*t*) is the firing rate of neuron *i, θ* = 0.01 is a target sparsity threshold (approximately 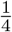 of the mean |*r*_*i*_(*t*) | observed in unregularized RNNs), and [·]^+^ denotes the rectifier linear unit function. Training employed the same procedures as for the CMM (Sec. 4.4), with a learning rate of 10^−4^ and an integration timestep *dt* = 5 ms to mitigate vanishing gradients [Sussillo et al., 2015].

#### Attempting to emulate CMM computational characteristics with RNNs

We tested whether the optimized rate-RNN control (Eqs. 14-18) could reproduce three CMM-like signatures when trained on the visual-change-detection and evidence-accumulation tasks: (1) condition-dependent sequential activity patterns (Sec. 4.5), (2) neural sparseness (Sec. 4.6), and (3) strong sensitivity to targeted silencing of high-information units. The hyperparameter search included:

1. Activation Regularization (*λ*_reg._): Varied logarithmically from 10^−8^ to 10^1^. Any *λ*_reg._ ≥ 10^−8^ substantially degraded task performance (test accuracy near chance). Population sparseness increased moderately (e.g., from 0.28 at *λ*_reg._ = 0 to 0.35 at *λ*_reg._ = 10^−3^ on visual-change-detection) but at the cost of task competency. Lifetime sparseness showed a non-monotonic relationship with *λ*_reg._ (e.g., *S*_*L*_ = 0.14 for *λ*_reg._ = 0; *S*_*L*_ = 0.17 for *λ*_reg._ = 10^−8^; *S*_*L*_ = 0.05 for *λ*_reg._ = 10^−4^). Given the primacy of task performance for meaningful comparison, *λ*_reg._ was set to 0 for final RNN evaluations unless noted.
2. Learning Rate: Explored logarithmically (10^−6^ to 10^−1^). A rate of 10^−4^ provided optimal convergence and accuracy. Variations in learning rate did not substantially alter the model’s sequential activity, sparseness, or functional segregation characteristics once task performance was saturated.
3. Activation function: Explored tanh, sigmoid, relu and silu. None changed the qualitative conclusion.
4. Weight Initialization: Compared standard Gaussian initialization 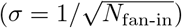 with initialization mirroring CMM’s. No significant differences in emulating CMM behaviors were found when task performance was matched. Standard Gaussian initialization was adopted.

Under these tested settings, the rate-RNN learned the task but did not reproduce CMM-like sequence condition dependence or CMM-like lifetime sparseness. These results should be interpreted as limitations of this matched rate-RNN control, not as a claim about all possible recurrent architectures.

#### Comparative statistical analysis

To assess the architectural contributions of spatially structured local connectivity and neuronal-type diversity, we organized four spiking models into a fully crossed 2 × 2 factorial design. Connectivity had two levels, V1-like and random; Diversity had two levels, full neuronal diversity and reduced excitatory/inhibitory types. The four cells were: CMM, with V1-like connectivity and full neuronal diversity; CMM1, with V1-like connectivity and reduced diversity; CMM2, with random connectivity and full neuronal diversity; and RSNN, with random connectivity and reduced diversity. The rate-based RNN was excluded from this factorial analysis because it used rate units rather than spiking neurons and therefore was not nested within the same experimental design.

Each factorial cell contained *N* = 10 independently trained model instances. The primary endpoints were defined before statistical testing as three temporal-order metrics and two sparseness metrics. For temporal-order metrics, we analyzed rank-order dissimilarity,

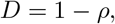

where *ρ* is the Spearman rank correlation between the peak-latency orders of two conditions. Larger values of *D* indicate stronger condition dependence of temporal ordering. The temporal-order metrics were change/no-change dissimilarity, current-image dissimilarity, and preceding-image dissimilarity. The sparseness metrics were lifetime sparseness and population sparseness.

For each metric, we fitted a two-way ANOVA model with interaction:

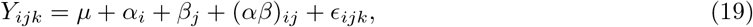

where *µ* is the grand mean, *α*_*i*_ is the main effect of Connectivity, *β*_*j*_ is the main effect of Diversity, (*αβ*)_*ij*_ is the Connectivity × Diversity interaction, and *ϵ*_*ijk*_ is the residual error for the *k*-th independently trained model instance in cell (*i, j*). Because the design was balanced and orthogonal, the factor tests were order-invariant.

For each effect, the *F*-statistic was computed as the ratio of the effect mean square to the residual mean square. Partial eta-squared was computed as

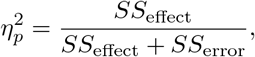

and omega-squared was computed as an adjusted effect-size estimate. Numerical consistency was verified by confirming that the reported *F*-statistics matched the values implied by 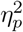 and the corresponding degrees of freedom.

Multiple-comparison correction was performed within two prespecified statistical families: temporal-order metrics and sparseness metrics. Within each family, the *P*-values for the Connectivity main effect, Diversity main effect, and Connectivity × Diversity interaction were corrected using the Holm procedure. We report *F*-statistics, Holm-corrected *P*-values, partial eta-squared 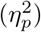, omega-squared (*ω*^2^), and 95% bootstrap confidence intervals for 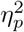. Bootstrap confidence intervals were obtained by resampling independently trained model instances with replacement within each factorial cell.

Simple-main-effects analyses were reserved for metrics in which the Connectivity × Diversity interaction remained significant after Holm correction. Because none of the interactions survived correction in the final analysis, simple-main-effects tests were not used for confirmatory interpretation.

### 4.8 Find early-informer neurons

Early-informer neurons are defined as those neurons whose mutual information between single neuron activity and the network decision are high. We select with the descending rank of mutual information. To calculate the mutual information, we binned the spike counts of each neuron into 10 uniformly distributed bins between the minimum and maximum spike count observed for that neuron within a window. To ensure the identification of early-informer neurons was not dependent on hyperparameter selection, windows with 10, 30, 50, and 70 ms were tested. The overlap of the top 200 neurons (early-informer neurons) remained above 92% across all schemes, demonstrating that this functional segregation is a robust property of the architecture and not an artifact of the chosen binning estimator.

We then established an empirical joint distribution for the binned spike count and the network decision and computed the mutual information using the below formula.

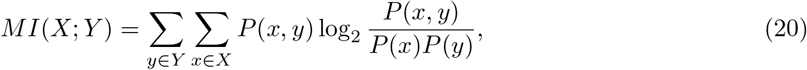

where *X* is the set of firing activities of each neuron within a 50 ms window, and *Y* is the set of network decisions (either change or no-change). *P* (*x, y*) is the joint probability distribution of spike count and network decision, while *P* (*x*) and *P* (*y*) are the marginal probability distributions of spike count and network decision, respectively.

To estimate *P* (*x, y*), we calculated the spike count of each neuron within a 50 ms window and established an empirical joint distribution by counting the number of occurrences of each possible combination of spike counts and network decisions across 100,000 trials. We then normalized the joint distribution to obtain a probability distribution. We estimated *P* (*x*) and *P* (*y*) in a similar way by counting the number of occurrences of each possible value of spike count and network decision, respectively, across all trials.

### 4.9 UMAP

We applied an exponential filter with a time constant of 20 ms to the spike output of each neuron for 8 new images that had not been used during training. We then discarded all but the 1,500 most important principal components of these network states (explain 38% of variance on visual-change-detection task and 41% of variance on evidence-accumulation task, as a compromise for the memory consumption), and embedded these into 2D space by UMAP. These projected network states were recorded for every ms, represented by a dot in Fig. 7.

UMAP (Uniform Manifold Approximation and Projection) is a nonlinear dimensionality reduction algorithm designed to maintain the inherent structure of high-dimensional data when projected into a lower-dimensional space [McInnes et al., 2018]. Unlike linear techniques, UMAP focuses on preserving the pairwise distances between neighboring data points. In other words, it strives to conserve the local relationships or similarities among points in the original high-dimensional data. In this study, UMAP was used to embed the network activity during task performance into 2D space. Specifically, we used the UMAP implementation from the Python library umap, with the following parameters: `n_neighbors` = 200, `min_dist` = 0.1, and `metric` = euclidean.

The resulting 2D embeddings represent the low-dimensional trajectories of the network states during image processing, and were recorded for every ms. Each point in the 2D space represents a network state at a given time point, and is displayed as a dot in Fig. 7. The trajectory of the network states can be visualized by connecting these dots in chronological order.

## Acknowledgments

The authors thank Sandra Diaz for advice and help in using supercomputers. This work was supported by the National Key R&D Program of China, Project Number 2025YFA1016700. This work was also supported by the National Natural Science Foundation of China (NSFC) under Grant 62576011. This research was partially supported by the Human Brain Project (Grant Agreement number 785907) of the European Union and a grant from Intel. Computations were carried out on the Human Brain Project PCP Pilot Systems at the Jülich Supercomputing Centre, which received co-funding from the European Union (Grant Agreement number 604102). This research was funded in part by the Austrian Science Fund (FWF) 10.55776/COE12.

## Supplementary Information

### 1 Supplementary Note 1: Tested recurrent controls show limited cortical-like temporal organization

A common approach in computational neuroscience is to model cortical function using artificial recurrent neural networks (RNNs) with rate units and random connectivity. Such networks can learn complex cognitive tasks without detailed biological architecture [Sussillo and Barak, 2013, Sussillo et al., 2015, Yang et al., 2019, Yang and Wang, 2020, Pollock and Jazayeri, 2020, Driscoll et al., 2024]. We therefore tested whether sparse condition-dependent temporal ordering would emerge in a matched rate-RNN control trained on the visual-change-detection task using BPTT (Methods Sec. 4.7). The RNN learned the task with high accuracy (> 90%), but its internal activity differed from the V1 and CMM signatures.

The RNN units tended to remain active over longer time windows, including periods outside the task-relevant response window. More importantly, the temporal order of activity was less sensitive to task condition than in V1 or the CMM. When the order estimated from change trials was applied to no-change trials, the middle panel remained similar to the condition-specific sorting, indicating weaker condition dependence of peak-latency order (Fig. 4).

Quantitatively, the RNN showed higher rank correlation across task conditions than mouse V1 or the CMM (Fig. 6A-C). It also showed lower lifetime sparseness than the CMM and mouse V1 (Fig. 6D,E). These results indicate that this tested rate-RNN control learned the task through a different internal strategy.

The same control also showed weaker sensitivity to targeted silencing of high-information units. To obtain a performance drop comparable to silencing 200 early-informer neurons in the CMM, a much larger number of RNN units had to be silenced (Fig. S3B).

Taken together, these analyses show that the particular rate-RNN control tested here does not reproduce the sparse, condition-dependent spike-order signature observed in the CMM and mouse V1. This should not be generalized to all recurrent architectures; modern adaptive, modular, low-rank, and gated recurrent models remain important baselines for future work.

### 2 Supplementary Note 2: Details to the evidence-accumulation task

Besides the visual-change-detection task, we trained another task in the CMM and variant control models: evidence-accumulation task (Fig. 1C, Sec. 4.2 in Methods) [Morcos and Harvey, 2016, Engelhard et al., 2019]. Much like the visual-change-detection task in the CMM, the evidence-accumulation task utilizes the temporal sequence of peak activity to perform computations (Fig. S1A). Note that in Fig. S1A, we used the sequence of left-right-left-left-left for the turning-left trials and the sequence of left-right-left-right-right for the turning-right trials, in order to better demonstrate the difference between two rank orders. The sequential patterns for turning left and turning right are distinctly different.

Using a similar method to calculate the mutual information between neural activity and network decision, we discovered that silencing a relatively small number of neurons (for instance, 100 with high mutual information: early-informer neurons) dramatically reduces task performance (Fig. S1B). We selected neurons to silence based on the descending order of their mutual information with the network decision calculated within a specific time period. We observed that neuronal activities carry increasingly higher information as they approach the response window, reflected by the severe reduction of task performance (Fig. S1B).

In addition, we calculate the rank correlation between trails of turning left and right, sparseness, and test accuracy in Fig. S2, suggesting similar conclusion as in visual-change-detection task. Also, we computed the mutual information between the neural activity over a temporal segment and the current state of accumulated cues (i.e., when the number of left cues is greater than, less than, or equal to the number of right cues). We found a similar effect when silencing neurons that exhibited high values of this type of mutual information.

To illustrate the computational progression of the CMM in the evidence-accumulation task, we employed UMAP to embed its activity vectors into two dimensions, mirroring the method used in the visual-change-detection task. For ease of visualization and analysis, we held the first three cues constant as left-right-left. The processing of each unique image generates two bundles of trajectories, with trial-to-trial variations arising from the sequence of cues and intrinsic network noise (Fig. 8A). These trajectories exhibit three nested condition-dependent separations in the embedding, providing a visualization of the computational progression of the trained network (see Fig. 8B). By silencing 100 early-informer neurons (neurons with the highest mutual information with the network decision in a certain window), these bifurcations can be flipped (Fig. 8C). This behavior mirrors that observed in the visual-change-detection task (Fig. 7).

**Figure S1:**
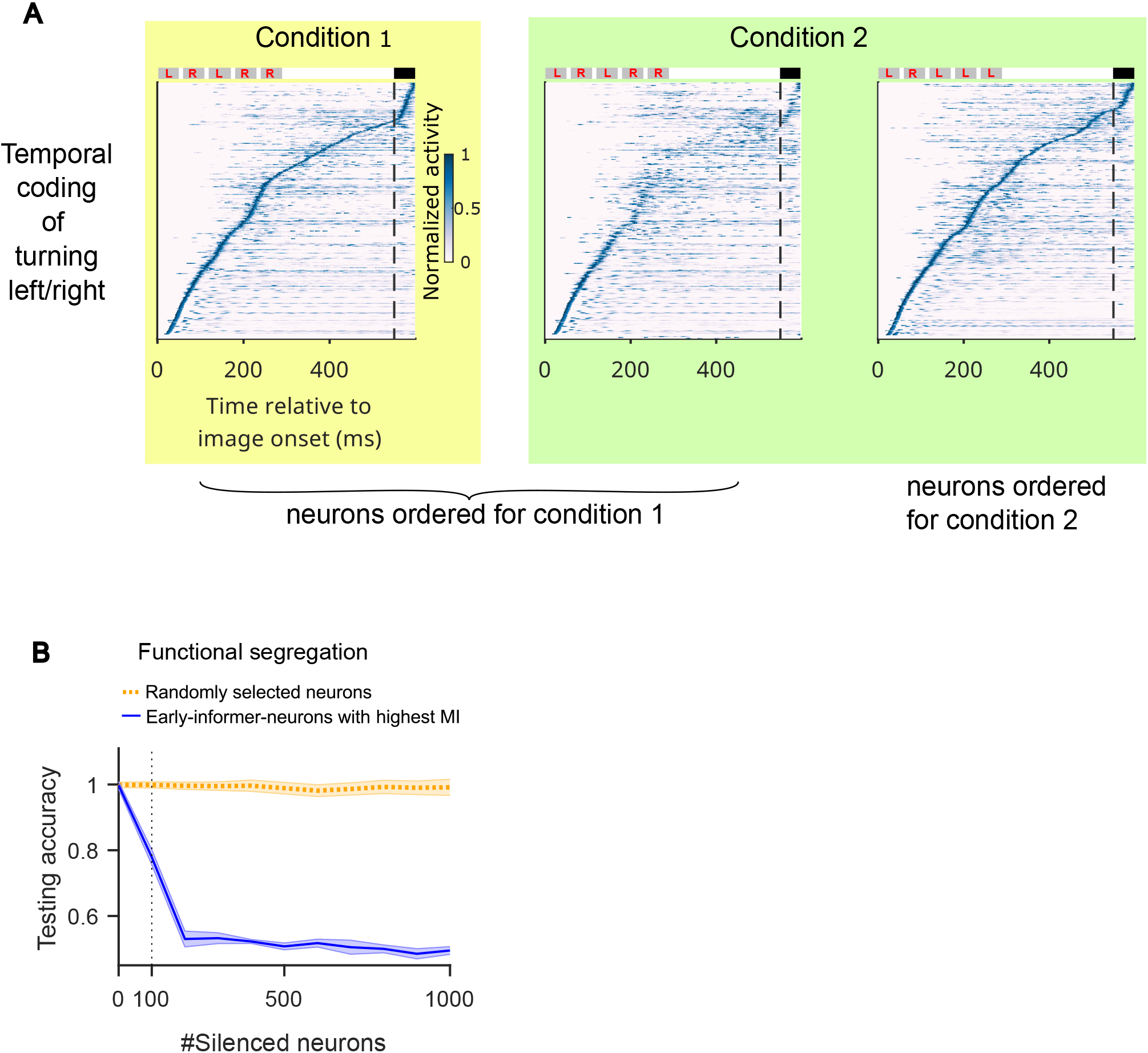
Temporal organization of computations for solving the evidence-accumulation task in the CMM. **(A)** Comparison of neural sequences between turning left (condition 1) and right (condition 2) trials. Left: Activity during right trials, with neurons sorted by their peak firing time in this condition, revealing a distinct temporal sequence. Middle: Activity during left trials, forced into the same neuronal order as the change condition on the left. The resulting blurred pattern demonstrates that the firing order is significantly different between the two conditions. Right: Activity during left trials, now sorted by its own optimal firing order, revealing its own unique and clear temporal sequence. **(B)** Causal impact of specific neurons on the network decision for this task: Task performance quickly decreases when early-informer neurons, ranked by their MI during the third cue presentation coinciding with the network decision, are silenced (represented by the blue curve). On the other hand, task performance is robust to silencing the same number of randomly selected neurons (dotted yellow curve). Both curves show average values for 10 CMMs where different random seeds were used. The shaded area represents the SEM across 10 models. Compare with the analogous plot in Fig. S4 B for the visual-change-detection task.

**Figure S2:**
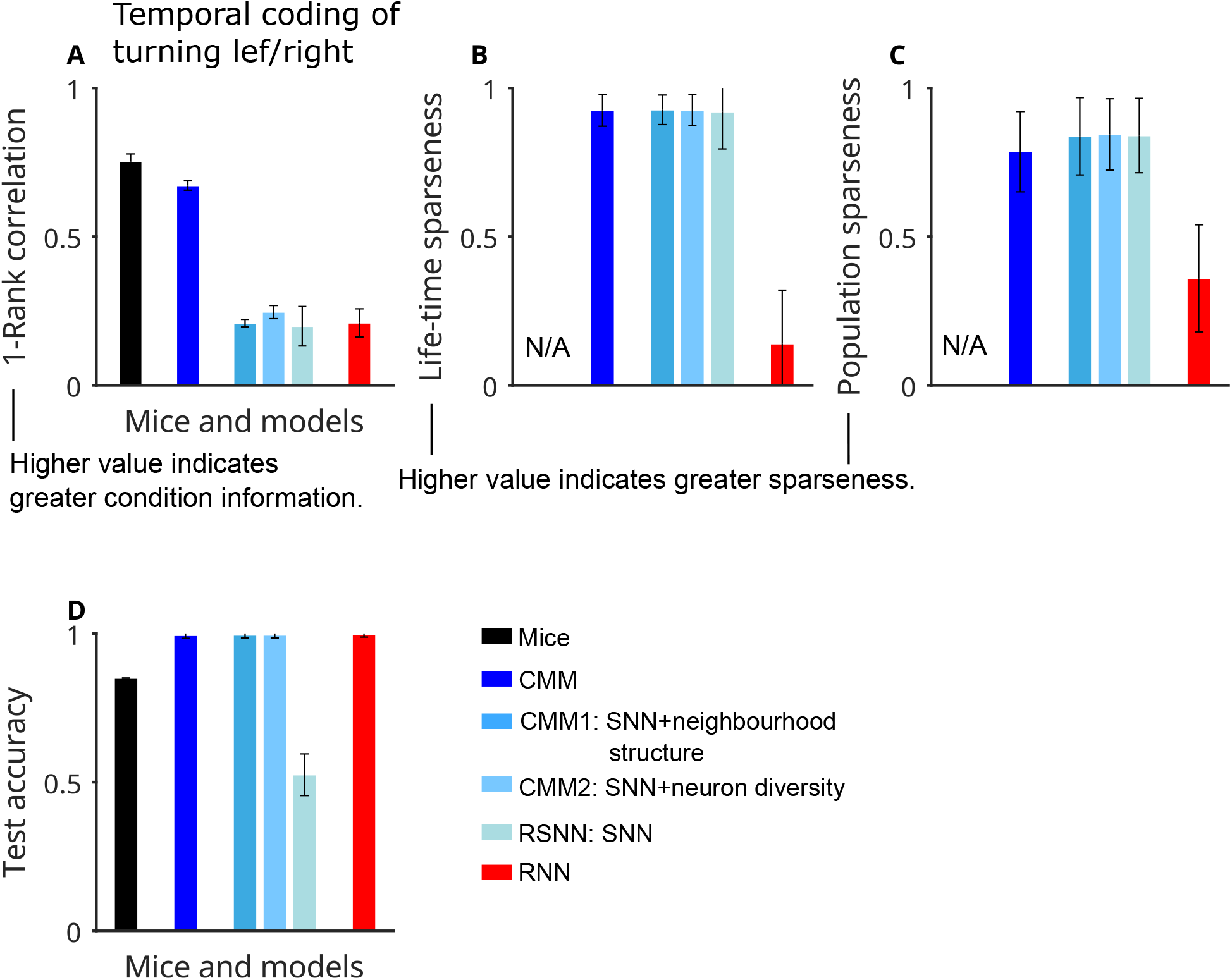
Temporal-coding information, sparseness and task performance in the evidence-accumulation task. **(A)** 1 - Spearman rank order correlation between change/no-change trial conditions. Larger values indicate greater condition-specific information in firing order. The value of mouse was calculated based on the V1 data provided in [Koay et al., 2022]. **(B)** Comparison of lifetime sparseness among models. **(C)** Same as in **(B)** but for population sparseness. **(D)** Comparison of test accuracy among models and mouse V1. The mouse performance was estimated from Fig. 1C of [Morcos and Harvey, 2016] when the number of cues was 6. We used 5 cues in our simulations. **(A-D)** share the same color scheme. Error bars are standard deviations calculated among 10 models.

**Figure S3:**
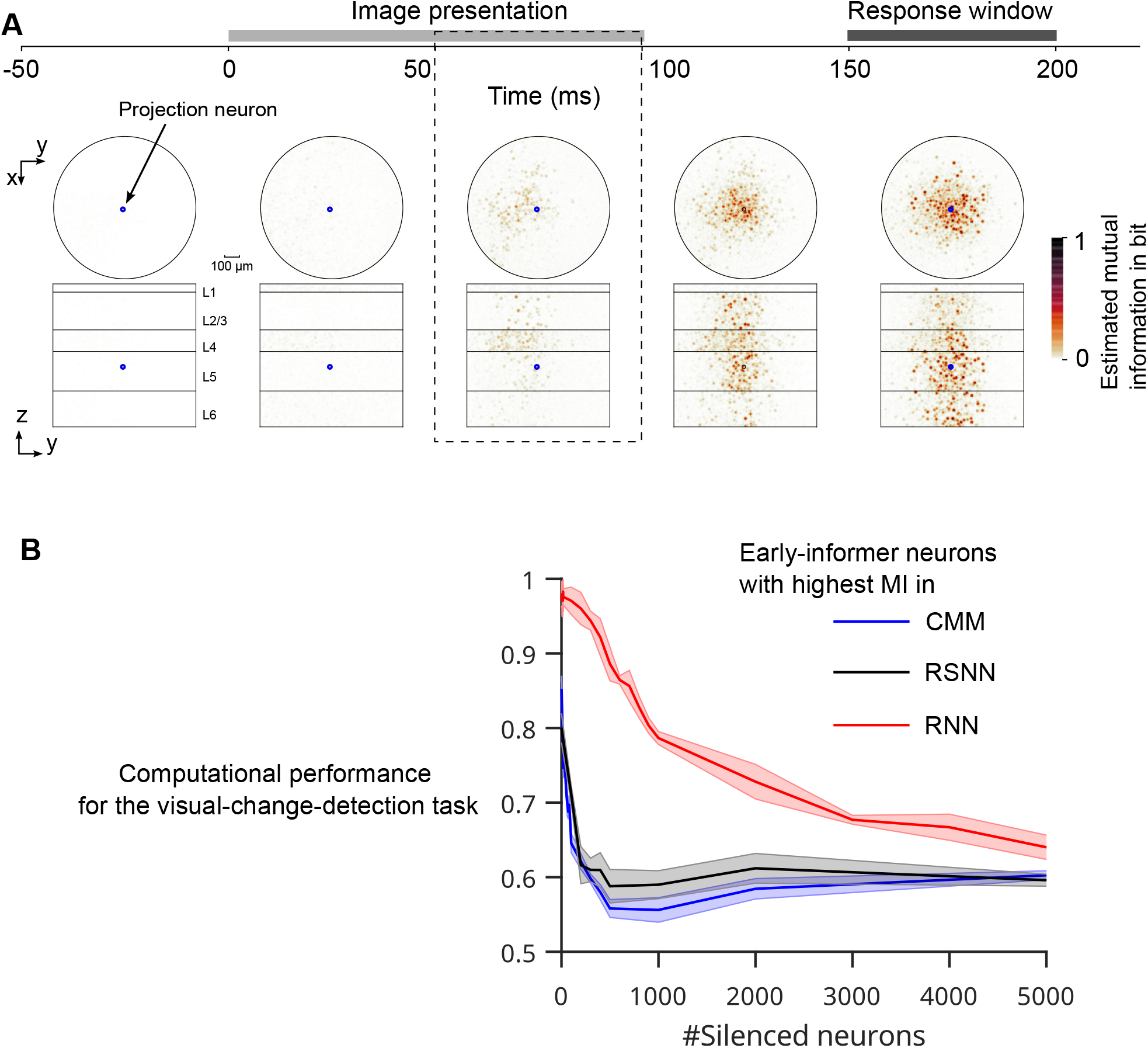
Functional segregation in the CMM for the visual-change-detection task. **(A)** The mutual information (MI) between activities in 50 ms windows of single neurons and the change/no-change decision of the network (as emerging during the subsequent response window) is estimated for each neuron. For neurons that overlap in the projection from 3D to 2D, the maximum value is visualized to avoid dark points arising from accumulation of small contributions from several neurons. The critical time period from 50 to 100 ms after image onset is marked by dashed lines; neurons with high mutual information in this period are called early-informer neurons. **(B)** Effect of targeted silencing on task performance for the CMM, RSNN, and RNN. In the CMM, silencing high-MI early-informer neurons identified during the [50, 100] ms interval reduces performance more strongly than random silencing. A similar vulnerability is observed in the RSNN, whereas the RNN is less sensitive to the loss of high-importance units. Curves show averages over 10 independently trained model instances; shaded areas denote SEM.

**Figure S4:**
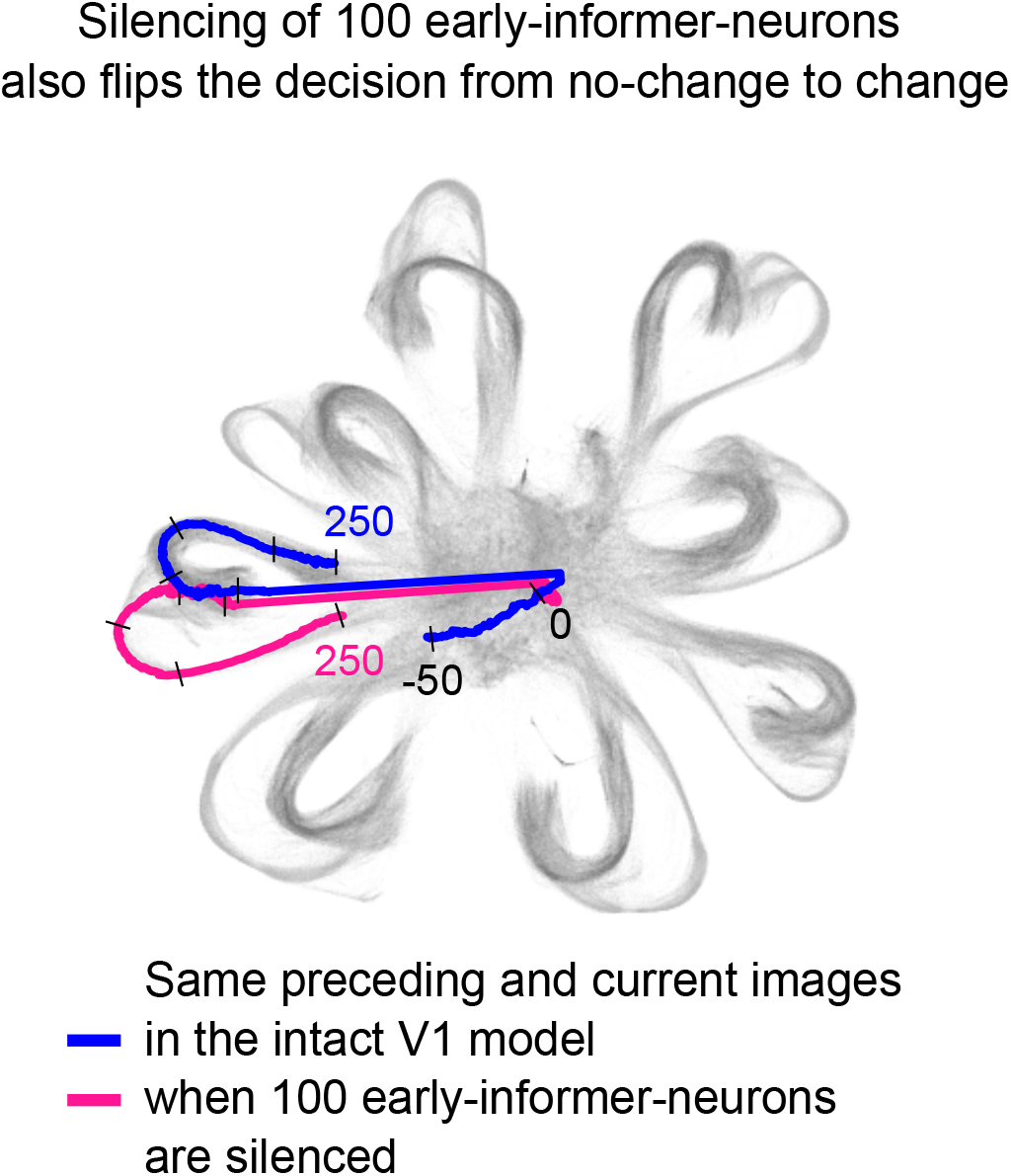
Silencing of 100 neurons can also flip a trajectory from the bundle for no-change to the change bundle of trajectories in the visual-change-detection task. In Fig. 7D, we demonstrated that silencing 100 early-informer neurons can cause the trajectory of network states to flip from the bundle for change to the bundle for no-change trials. We show here that silencing of the same 100 neurons can also flip the network bifurcation in the other direction.

**Table S1:** Two-way factorial ANOVA for architectural effects on temporal-order dissimilarity and sparseness.

| Metric | Connectivity | Diversity | Connectivity × Diversity |
| --- | --- | --- | --- |
| *Temporal-order dissimilarity, $D = 1 - \rho$* |  |  |  |
| Change/no-change | $F_{1,36} = 61.74$<br>$P_{\text{Holm}} = 2.30 \times 10^{-8}$<br>$\eta_p^2 = .632$ [.395, .863] | $F_{1,36} = 17.16$<br>$P_{\text{Holm}} = 7.93 \times 10^{-4}$<br>$\eta_p^2 = .323$ [.091, .685] | $F_{1,36} = 5.90$<br>$P_{\text{Holm}} = .051$<br>$\eta_p^2 = .141$ [.005, .516] |
| Current image | $F_{1,36} = 28.71$<br>$P_{\text{Holm}} = 3.50 \times 10^{-5}$<br>$\eta_p^2 = .444$ [.258, .714] | $F_{1,36} = 20.26$<br>$P_{\text{Holm}} = 3.41 \times 10^{-4}$<br>$\eta_p^2 = .360$ [.159, .655] | $F_{1,36} = 6.26$<br>$P_{\text{Holm}} = .051$<br>$\eta_p^2 = .148$ [.009, .456] |
| Preceding image | $F_{1,36} = 28.55$<br>$P_{\text{Holm}} = 3.50 \times 10^{-5}$<br>$\eta_p^2 = .442$ [.291, .684] | $F_{1,36} = 32.69$<br>$P_{\text{Holm}} = 1.33 \times 10^{-5}$<br>$\eta_p^2 = .476$ [.321, .716] | $F_{1,36} = 1.33$<br>$P_{\text{Holm}} = .257$<br>$\eta_p^2 = .036$ [.000, .249] |
| *Sparseness* |  |  |  |
| Lifetime sparseness | $F_{1,36} = 1.19$<br>$P_{\text{Holm}} = 1.000$<br>$\eta_p^2 = .032$ [.000, .205] | $F_{1,36} = 1.19$<br>$P_{\text{Holm}} = 1.000$<br>$\eta_p^2 = .032$ [.000, .215] | $F_{1,36} = 1.19$<br>$P_{\text{Holm}} = 1.000$<br>$\eta_p^2 = .032$ [.000, .198] |
| Population sparseness | $F_{1,36} = 0.05$<br>$P_{\text{Holm}} = 1.000$<br>$\eta_p^2 = .001$ [.000, .136] | $F_{1,36} = 0.76$<br>$P_{\text{Holm}} = 1.000$<br>$\eta_p^2 = .021$ [.000, .202] | $F_{1,36} = 3.06$<br>$P_{\text{Holm}} = .534$<br>$\eta_p^2 = .078$ [.001, .312] |

Note. Temporal-order metrics were analyzed as rank-order dissimilarity $D = 1 - \rho$ , where larger values indicate stronger condition dependence of peak-latency order. $P$ -values were corrected within prespecified statistical families using the Holm procedure. Brackets indicate 95% bootstrap confidence intervals for partial eta-squared. Raw $P$ -values and $\omega^2$ values are provided in Source Data.

